# Loss of myeloid cannabinoid CB1 receptor confers atheroprotection by reducing macrophage proliferation and immunometabolic reprogramming

**DOI:** 10.1101/2023.04.06.535832

**Authors:** Yong Wang, Guo Li, Bingni Chen, George Shakir, Mario Volz, Emiel P.C. van der Vorst, Sanne L. Maas, Carolin Muley, Alexander Bartelt, Zhaolong Li, Nadja Sachs, Lars Maegdefessel, Maliheh Nazari Jahantigh, Michael Hristov, Michael Lacy, Beat Lutz, Christian Weber, Stephan Herzig, Raquel Guillamat Prats, Sabine Steffens

## Abstract

Although the cannabinoid CB1 receptor has been implicated in atherosclerosis, its cell-specific effects in this disease are not well understood. Here, we report that male mice with myeloid-specific *Cnr1* deficiency on atherogenic background developed smaller lesions and necrotic cores than controls, while only minor genotype differences were observed in females. Male *Cnr1* deficient mice showed reduced arterial monocyte recruitment and macrophage proliferation with less inflammatory phenotype. The sex-specific differences were reproducible in bone marrow derived macrophages and blunted by estradiol. Kinase activity profiling revealed a CB1-dependent regulation of p53 and cyclin-dependent kinases. Transcriptomic profiling further unveiled chromatin modifications, mRNA processing and mitochondrial respiration among the key processes affected by CB1 signaling, which was supported by metabolic flux assays. Chronic administration of the peripherally-restricted CB1 antagonist JD5037 inhibited plaque progression and macrophage proliferation, but only in male mice. Finally, *CNR1* expression was detectable in human carotid endarterectomy plaques and inversely correlated with proliferation, oxidative metabolism and inflammatory markers, hinting to a possible implication of CB1-dependent regulation in human pathophysiology. In conclusion, impaired CB1 signaling in macrophages is atheroprotective by limiting their arterial recruitment, proliferation and inflammatory reprogramming. The importance of macrophage CB1 signaling seems to be more pronounced in male mice.

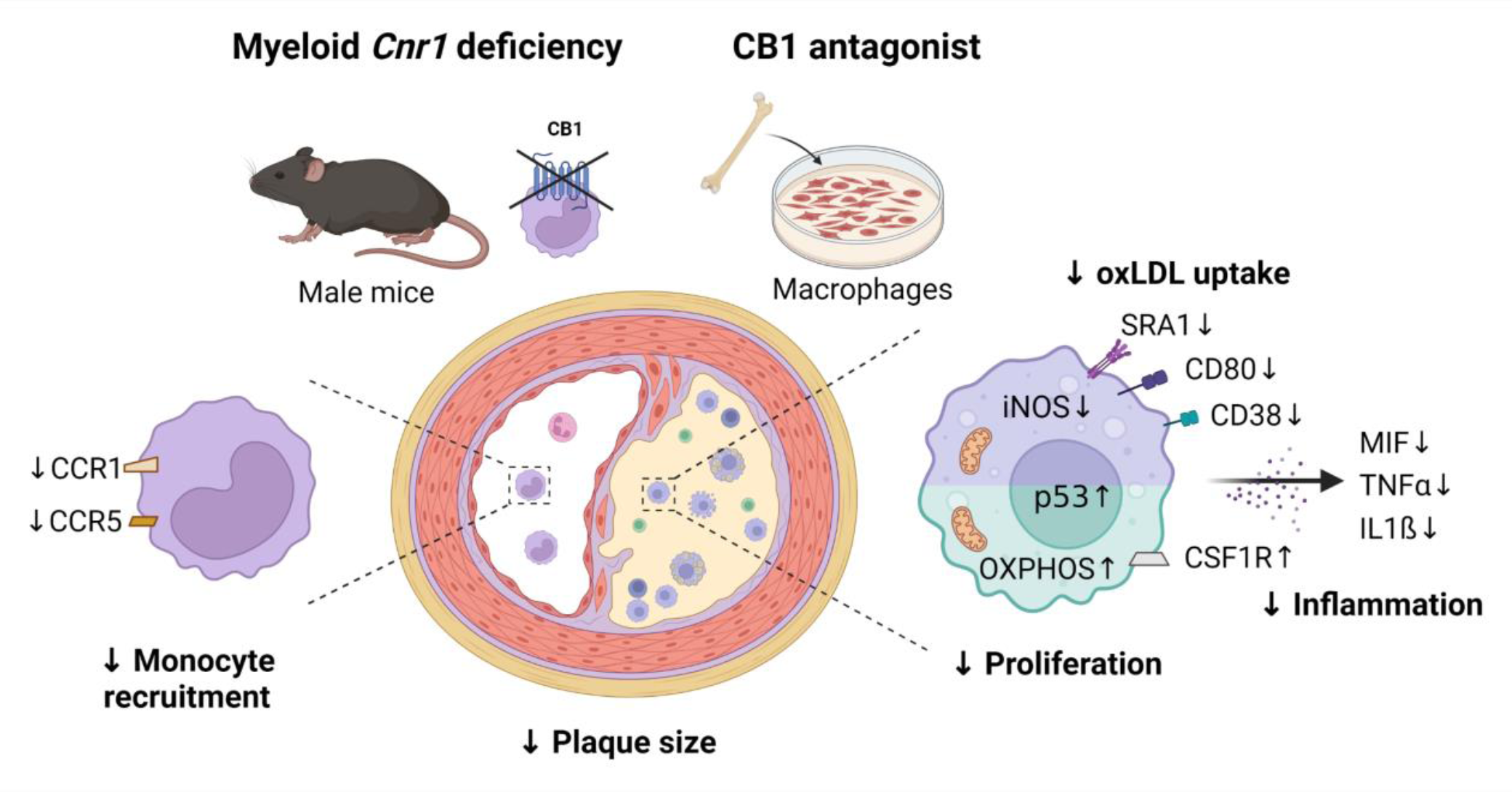

**Graphical summary** (created with BioRender.com)

## Introduction

Cardiovascular diseases (CVD) are the primary cause of death worldwide. While there have been significant advancements in lipid-lowering therapies and positive results from initial clinical studies targeting inflammation in CVD patients, atherosclerosis remains a significant health burden. This is due to an increasing life expectancy and ageing population in our society, as well as an increased prevalence of lifestyle-related cardiometabolic risk factors. Thus, improving CVD prevention and therapy remains a central question in biomedical research with the aim to achieve a more efficient and reliable identification of new targets for translation to drug development. In particular, the development of novel therapeutics that block atherosclerosis-specific inflammatory pathways without immunosuppressive side effects remains an unmet need to optimize immunotherapy for CVD (1).

Endocannabinoids are a group of arachidonic acid-derived lipid mediators with pleiotropic cardiometabolic and immunomodulatory properties. Circulating endocannabinoid levels are elevated in patients with coronary artery disease (2), myocardial infarction (3), chronic heart failure (4), and obese patients with coronary endothelial dysfunction (5). In a mouse model of atherosclerosis, cannabinoid CB1 receptor antagonism with rimonabant inhibited plaque formation (6). Other investigators failed to observe effects of rimonabant on plaque size but found improved aortic endothelium-dependent vasodilation as well as decreased aortic ROS production and NADPH oxidase activity (7). More recently, the soybean isoflavone genistein was identified as a CB1 antagonist through in silico screening and was reported to inhibit plant cannabinoid-mediated inflammation and oxidative stress in endothelial cells and plaque progression in a mouse model (8).

Peripherally-restricted CB1 antagonists, which are devoid of undesirable neuronal actions in the brain, improved metabolic dysfunction in experimental mouse models (9–11). Liver-specific CB1 deficiency blunted glucocorticoid-induced dyslipidemia, but not the obesity phenotype (9). Inducible adipocyte-specific *Cnr1* deficiency was sufficient to protect adult mice from diet-induced obesity and associated metabolic alterations and reversed the phenotype in obese mice (12). In pancreas-infiltrating macrophages, CB1 signaling exhibited pro-inflammatory effects through activation of the NLRP3 inflammasome complex, thereby contributing to beta-cell loss in experimental type 2 diabetes (13). The experimental *in vivo* evidence in this study was based on combined treatment with peripherally active CB1 antagonists and clodrononate liposomes for macrophage depletion or siRNA-mediated CB1 knockdown, but not cell-specific genetic *Cnr1* depletion. In particular, CB1 siRNA treatment prevented hyperglycemia, macrophage infiltration, and *Nlrp3* expression, similar to the effects observed by JD5037 or clodronate treatment.

In atherosclerosis, monocyte-derived macrophages are the main leukocyte subset recruited to the activated endothelium and accumulating within plaques. *In vitro* data point to a possible regulation of macrophage cholesterol metabolism by cannabinoids (14), although the *in vivo* relevance of this mechanism in plaque macrophages is still unexplored. The proposed role of CB1 in macrophage cholesterol metabolism and inflammatory signaling, as reported in experimental diabetes, suggest a crucial role for CB1 in regulating the macrophage phenotype in the atherosclerotic plaque. However, studies based on cell-targeted genetic disruption of CB1 in macrophages in experimental atherosclerosis are still lacking.

To address this issue, we generated a mouse model with myeloid cell-specific *Cnr1* deficiency to study the effects of myeloid CB1 signaling in early and advanced stages of atherosclerosis. Furthermore, we investigated the therapeutic benefit of peripheral CB1 antagonism for inhibiting plaque progression.

## Results

*Inactivation of myeloid CB1 enhances early and advanced atherosclerotic plaque formation* We first confirmed the expression of the CB1 encoding gene *Cnr1* in sorted mouse blood leukocytes using droplet digital PCR, revealing highest mRNA levels in lymphocytes, and slightly lower levels in monocytes and neutrophils (Supplemental Figure 1A). In bone marrow-derived macrophages (BMDMs) isolated from *Apoe^-/-^* mice, inflammatory stimulation with LPS or oxLDL increased *Cnr1* expression levels (Supplemental Figure 1B), which is in line with previous findings in macrophage cell lines (2, 14). We subsequently crossbred *Cnr1^flox/flox^* mice with *LysM^Cre^* mice, using the apolipoprotein E deficiency (*Apoe^-/-^*) background as a model of atherosclerosis. *In situ* hybridization combined with immunofluorescence staining confirmed *Cnr1* expression by CD68+ plaque macrophages in aortic root sections of control mice (*Apoe^- /-^LysM^Cre^*), while the *Cnr1* signal was undetectable in plaques of mice with myeloid *Cnr1* deficiency (*Apoe^-/-^LysM^Cre^Cnr1^flox/flox^*; Supplemental Figure 1C). In cultured bone marrow-derived macrophages (BMDMs) obtained from *Apoe^-/-^LysM^Cre^Cnr1^flox/flox^*mice, qPCR analysis revealed a 50% lower *Cnr1* expression level compared to *Apoe^-/-^LysM^Cre^* controls (Supplemental Figure 1D). This is in accordance with the use of heterozygous *LysM^Cre^* mice throughout our entire study, to avoid a full disruption of *LysM* expression due to the *Cre* insertion into the first coding ATG of the lysozyme M gene (15).

After successful establishment of the mouse model, we asked whether depletion of CB1 in myeloid cells affected early stages of atherosclerotic plaque formation. For this purpose, we initially used two separate control groups, in order to exclude any effect of the *Cnr1^flox/flox^*or *LysM^Cre^* transgene insertion on the atherosclerosis phenotype, independent of the myeloid *Cnr1* depletion. In subsequent experiments, *Apoe^-/-^LysM^Cre^* mice were consistently used as controls. After 4 weeks of Western diet (WD) feeding, male mice with myeloid *Cnr1* deficiency had smaller plaques within the aortic roots compared to the *Apoe^-/-^Cnr1^flox/flox^* or *Apoe^-/-^LysM^Cre^* control group (Figure 1A-D). Conversely, no effect of myeloid *Cnr1* deficiency on early atherogenesis was observed in female mice. The smaller plaque size in males was linked to lower arterial macrophage accumulation, which was unaffected in females; nonetheless, both male and female plaques had a lower content of inflammatory iNOS^+^CD68^+^ macrophages (Figure 1E-G). However, myeloid *Cnr1* deficiency did not affect plaque neutrophil counts (Supplemental Figure 2).

**Figure 1.**
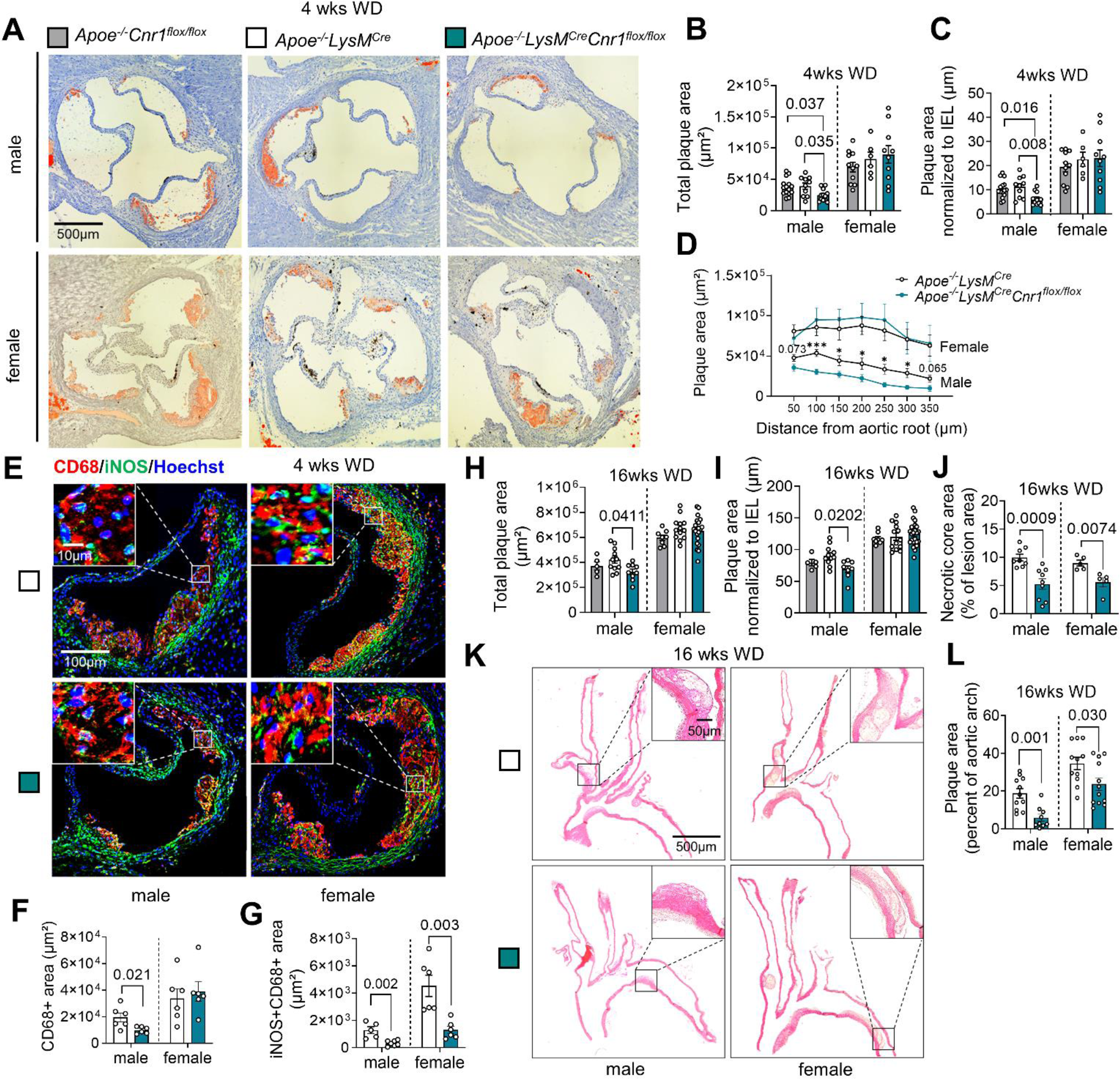
Impact of myeloid *Cnr1* deficiency on atherogenesis and plaque progression. **(A-D)** Representative Oil-Red-O (ORO) stains of aortic roots of male (n=12-15) and female (n=6-11) *Apoe^-/-^Cnr1^flox/flox^, Apoe^-/-^LysM^Cre^* and *Apoe^-/-^LysM^Cre^Cnr1^flox/flox^* mice after 4 weeks of Western diet (WD); Scale bar: 500 μm. (**B**) Quantification of absolute lesion area or (**C**) normalized to IEL. (**D**) Plaque area per aortic root section. (**E**) Double immunostaining of iNOS (green) and CD68 (red) in aortic root lesions of *Apoe^-/-^LysM^Cre^*and *Apoe^-/-^LysM^Cre^Cnr1^flox/flox^* mice after 4 weeks of WD. Nuclei were counterstained with Hoechst 33342 (blue). Scale bar: 10 μm (top) and 100 µm (bottom). (**F**) Quantification of lesional macrophages and (**G**) inflammatory macrophages identified by combined positive CD68 and iNOS staining (n=5–6). (**H**) Quantification of absolute lesion area or (**I**) normalized to IEL in aortic root sections after 16 weeks of WD (n=5–21). (**J**) Necrotic core area quantified in Masson Trichrome-stained aortic root sections after 16 weeks of WD (n=5–9). (**K**-**L**) Representative HE stains of aortic arches and plaque area quantification in male and female *Apoe^-/-^LysM^Cre^* and *Apoe^-/-^ LysM^Cre^Cnr1^flox/flox^* mice after 16 weeks of WD (n=9–12); Scale bar: 500 μm. Each dot represents one biologically independent mouse sample and all data are expressed as mean ± s.e.m. Two-sided unpaired Student’s t-test (**D, F, G, J** and **L**), one-way ANOVA followed by Tukey test (**B** and **C**) or Dunnett T3 test (**H** and **I**) were used to determine the significant differences. Male and female were analyzed independently (**B-D, F-J** and **L**). Exact p-values are shown or indicated as ***p<0.001 and *p<0.05 in (**D**).

At advanced plaque stage after 16 weeks WD feeding, a lower aortic root plaque size was again only found in male *Apoe^-/-^LysM^Cre^Cnr1^flox/flox^*mice when compared to *Apoe^-/-^LysM^Cre^* controls, with a similar phenotype observed in descending aortas (Figure 1H-I and Supplemental Figure 3). However, the necrotic core size was smaller in both male and female aortic sinus plaques while other plaque components were comparable between genotypes, indicative of a more stable plaque phenotype in mice lacking CB1 (Figure 1J and Supplementary Figure 4). Furthermore, the relative plaque area within aortic arches was smaller in both male and female *Apoe^-/-^LysM^Cre^Cnr1^flox/flox^*mice (Figure 1K-L). Depleting myeloid CB1 signaling also affected metabolic parameters upon long-term 16 weeks WD feeding, which were only significant in male mice and manifested as lower body weight gain and lower plasma cholesterol levels (Supplemental Figure 5). Together, these findings indicate that the effect of myeloid *Cnr1* deficiency on the plaque phenotype appears to be stage- and vessel-dependent and affected by the biological sex.

### Lack of myeloid CB1 reduces chemokine receptor expression and arterial monocyte recruitment

Monocyte recruitment from the blood is the major source for plaque macrophages, in particular during atherogenesis (16). We therefore asked whether myeloid *Cnr1* deficiency affected circulating myeloid counts. We found higher levels of circulating monocytes in both male and female mice with myeloid *Cnr1* deficiency after 4 weeks WD (Figure 2A-B), while neutrophil blood counts at this time point were only elevated in male, but not female *Apoe^-/-^ LysM^Cre^Cnr1^flox/flox^*mice (Supplemental Figure 6A-B). Remarkably, an inverted pattern was observed at advanced stage of ≥ 12 weeks WD, with lower monocyte and neutrophil counts in *Apoe^-/-^LysM^Cre^Cnr1^flox/flox^* mice compared to *Apoe^-/-^LysM^Cre^* controls (Figure 2A-B, Supplemental Figure 6A-B). No significant differences in myeloid cell counts were observed in the bone marrow between the genotypes, neither at baseline nor in WD condition, except for a moderate difference in neutrophil counts of female at early and advanced stage (Supplemental Figure 6C-F). Furthermore, there was no difference in lymphocyte blood counts between the genotypes (Supplemental Figure 6G-H). Based on these findings we hypothesized that the lack of CB1 in myeloid cells affects their arterial recruitment and possibly their life span.

**Figure 2.**
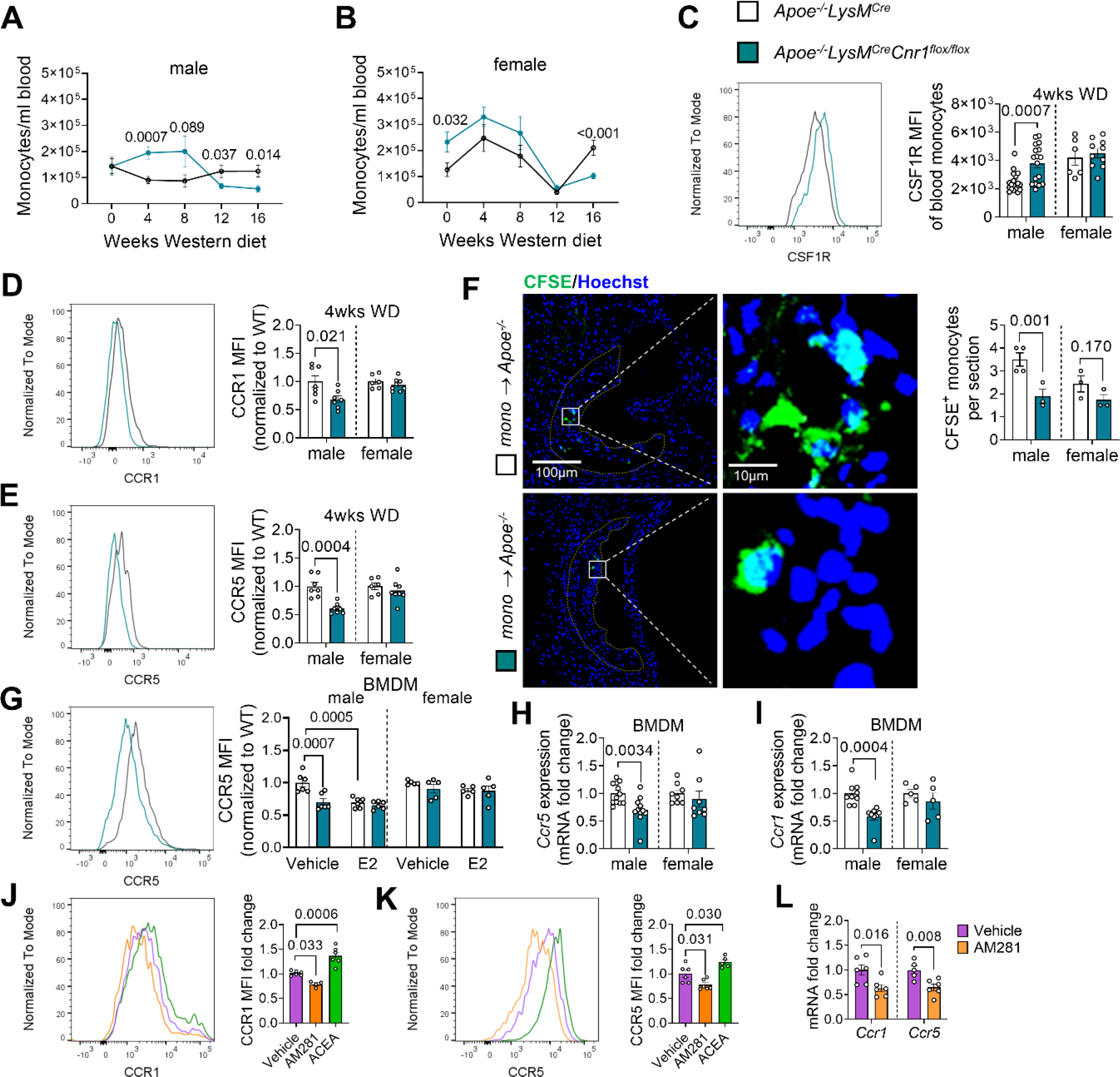
Impact of myeloid *Cnr1* deficiency on circulating leukocyte counts, chemokine receptor expression and arterial recruitment. **(A, B)** Number of circulating monocytes in male (7–19) and female (5–23) *Apoe^-/-^LysM^Cre^*and *Apoe^-/-^LysM^Cre^Cnr1^flox/flox^* mice assessed by flow cytometry. (**C**) Flow cytometric analysis of colony stimulating factor 1 receptor (CSF1R) expression on circulating monocytes after 4 weeks of WD (n=6–18). (**D**, **E**) Flow cytometric analysis of CCR1 and CCR5 surface expression on circulating monocytes after 4 weeks of WD (n=6–8). (**F**) Microscopy images and quantification of CFSE-labeled monocytes (green) recruited into aortic root lesions 48 h after injection into male *Apoe^-/-^* mice on WD for 4 weeks (n=3-4), Nuclei were stained with Hoechst 33342 (blue); Scale bar: 100 μm (left) and 10 µm (right). (**G**) Flow cytometric analysis of BMDM treated with vehicle or 10 µM estradiol for 24 h (n=3-6). (**H**, **I**) Gene expression levels in untreated BMDMs (n=5-12). (**J**, **K)** Flow cytometric analysis of BMDM from *Apoe^-/-^* mice treated with vehicle, 10 µM AM281 or 1 µM ACEA for 24 h (n=4-6). (**L**) Gene expression levels in *Apoe^-/-^* BMDMs treated with vehicle or 10 µM AM281 for 8 h (n=5-6). Each dot represents one biologically independent mouse sample and all data are expressed as mean ± s.e.m. Two-sided unpaired Student’s t-test (**A-F, H-I and L**), one-way ANOVA followed by Tukey test (**J** and **K**), two-way ANOVA followed by Tukey test (**G**). Male and female were analyzed independently (**A-I**).

In support of a pro-survival effect on monocytes, we noted that circulating male *Cnr1*-deficient monocytes had significantly higher surface levels of CD115, also known as colony-stimulating factor-1 receptor (CSF-1R) or macrophage-colony-stimulating factor receptor (M-CSFR; Figure 2C) (17, 18). Since plaque monocyte recruitment depends on the chemokine receptors CCR1 and CCR5 (19), we analyzed their surface expression on circulating monocytes and observed significantly lower expression levels in the blood of male *Apoe^-/-^LysM^Cre^Cnr1^flox/flox^* compared to *Apoe^-/-^LysM^Cre^* mice, while no difference was found in females (Figure 2D-E). The surface levels of CXCR2 on neutrophils were comparable between the genotypes, suggesting that neutrophil recruitment is unaffected by myeloid *Cnr1* deficiency (Supplemental Figure 6I). To strengthen our hypothesis that myeloid *Cnr1* deficiency leads to lower arterial monocyte recruitment, we performed an adoptive transfer experiment. Fluorescently labeled monocytes isolated from *Apoe^-/-^LysM^Cre^Cnr1^flox/flox^*or *Apoe^-/-^LysM^Cre^* mice were injected into *Apoe^-/-^*recipients pretreated with WD for 4 weeks. Fluorescence microscopic analysis of male aortic root sections transplanted with male monocytes revealed a significantly lower number of plaque-infiltrated *Cnr1*-deficient versus *Cnr1*-expressing monocytes, while the values in females did not reach significance (Figure 2F).

To study the CB1-regulated expression of CCR1 and CCR5 expression in more detail, we moved to *in vitro* experiments with BMDMs isolated from *Apoe^-/-^LysM^Cre^Cnr1^flox/flox^*or *Apoe^-/-^ LysM^Cre^* mice. A lower CCR5 surface expression was reproducible in male, but not female *Cnr1-*deficient BMDMs (Figure 2G). The potent estrogen receptor ligand estradiol suppressed the CCR5 surface expression in male *Apoe^-/-^LysM^Cre^*BMDMs, thereby blunting the difference between the control and *Cnr1* deficient group (Figure 2G). The qPCR analysis further revealed lower *Ccr1* and *Ccr5* mRNA levels in male *Cnr1-*deficient BMDMs (Figure 2H-I). Treatment of male *Apoe^-/-^* BMDMs with the selective synthetic CB1 agonist ACEA increased the CCR1 and CCR5 surface levels, whereas treatment with the CB1 antagonist AM281 led to lower surface levels compared to vehicle treated BMDMs (Figure 2J-K). Likewise, CB1 antagonism in male *Apoe^-/-^* BMDMs led to lower *Ccr1* and *Ccr5* mRNA levels (Figure 2L), which altogether suggests a direct regulatory effect of CB1 signaling on these chemokine receptors, possibly at the transcriptional or post-transcriptional level.

### Lack of myeloid CB1 inhibits macrophage proliferation via regulation of p53 signaling

Not only *de novo* recruitment, but also local proliferation determines the content of macrophages in the lesion (20). Co-immunostaining of macrophages and proliferation marker Ki67 revealed fewer proliferating macrophages per section in aortic roots of male *Apoe^-/-^ LysM^Cre^Cnr1^flox/flox^*compared to *Apoe^-/-^LysM^Cre^* mice (Figure 3A-B). Again, no difference was detected between the genotypes in females, while the proliferation rate was generally lower in female versus male plaque macrophages (Figure 3A-B). The same pattern was reproduced *in vitro* in male and female BMDMs, confirming lower proliferation rates in male *Apoe^-/-^ LysM^Cre^Cnr1^flox/flox^*compared to *Apoe^-/-^LysM^Cre^* macrophages (Figure 3C-D). This effect was blunted when treating male BMDMs with estradiol (Figure 3E). Likewise, the treatment of male, but not female *Apoe^-/-^* BMDMs with the CB1 antagonist AM281 inhibited the proliferation rate (Figure 3F). Conversely, treatment with the CB1 agonist ACEA increased the proliferation of male, but not female *Apoe^-/-^* BMDMs (Figure 3F). To address the underlying signaling pathways involved in the CB1-dependent regulation of cell proliferation, we performed a chip-based kinase activity profiling in BMDMs. Stimulating male *Apoe^-/-^*BMDMs with the CB1 agonist ACEA revealed a significant down-regulation of the activity of several kinases linked to p53 signaling and cyclin-dependent regulation of cell cycle (Figure 3G and Supplemental Figure 7A-B). The inhibition of male *Apoe^-/-^* BMDM proliferation by CB1 antagonism was not observed when pretreating the cells with the p53 inhibitor pifithrin-α (PFTα) (Figure 3H). Moreover, CB1 antagonism with AM281 resulted in higher nuclear translocation of phosphorylated p53, whereas the CB1 agonist ACEA lowered the levels of nuclear p53 compared to vehicle (Figure 3I and Supplemental Figure 7C).

**Figure 3.**
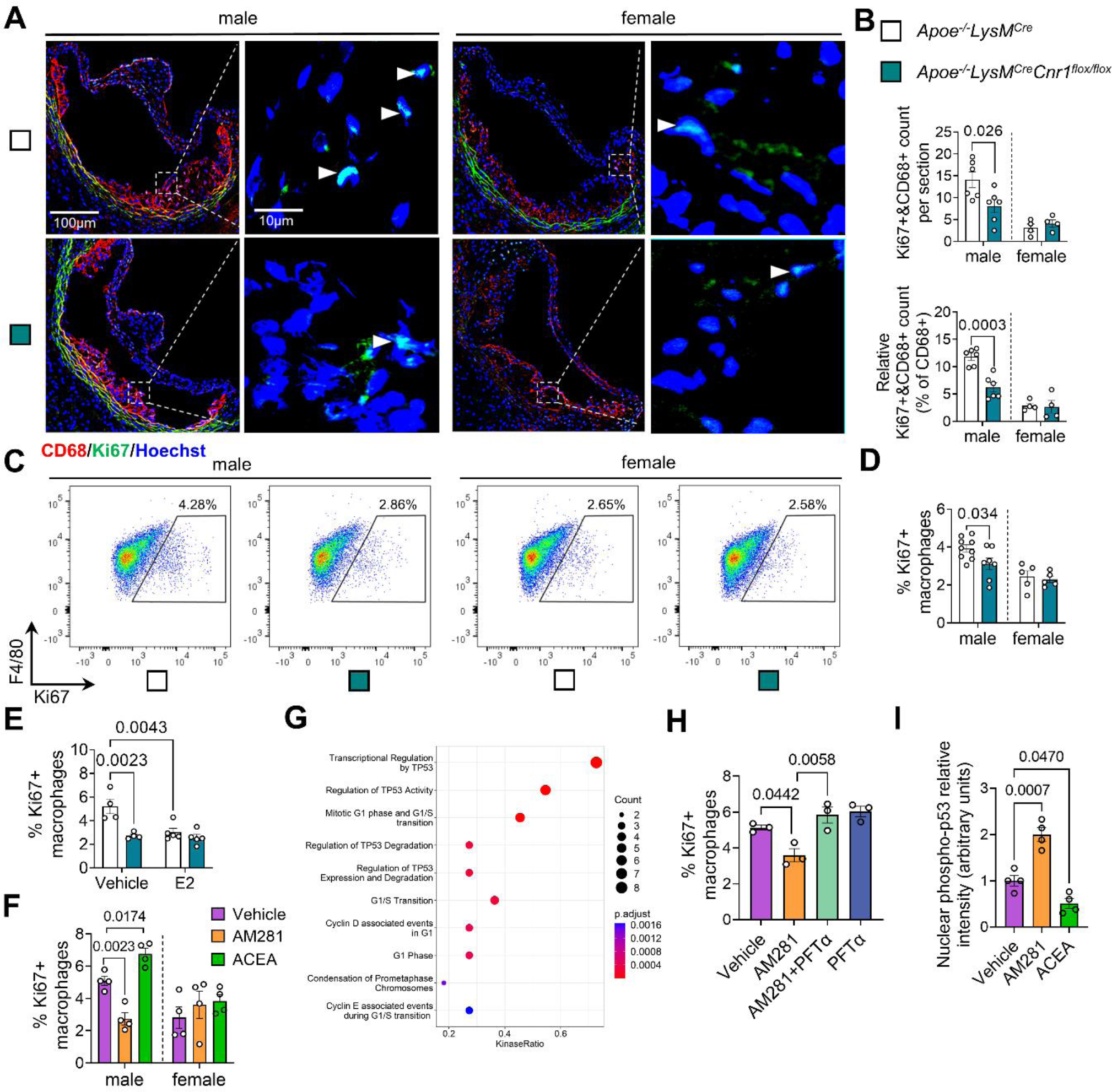
Role of CB1 and biological sex in macrophage proliferation. **(A)** Representative images of proliferating (Ki67+, green and marked with white arrowheads) macrophages (CD68+, red) in aortic root plaques after 4 weeks Western diet (WD); nuclei were counterstained with Hoechst 33342 (blue). scale bar: 100 μm (left) and 10 µm (right). (**B**) Total and relative counts of proliferating plaque macrophages (n=4-6). (**C**, **D**) Flow cytometric analysis of proliferation rates in untreated male and female BMDM isolated from *Apoe^-/-^LysM^Cre^*and *Apoe^-/-^LysM^Cre^Cnr1^flox/flox^* mice (n=5-9). (**E**) Proliferation rates in male BMDM treated with vehicle or 10 µM estradiol for 24 h (n=4-5). (**F**) Proliferating *Apoe^-/-^* BMDM treated with vehicle, 10 µM AM281 or 1 µM ACEA for 24 h (n=4). (**G**) Top 10 enriched kinase pathways in ACEA-stimulated male derived BMDMs (n=4). (**H**) Proliferation rates in BMDM treated with vehicle, AM281 (10 µM) and p53 inhibitor (PFT-α; 25 µM) for 24 h (n=3). (**I**) Nuclear transclocation of phosphorylated p53 in BMDM treated for 1 h with vehicle, AM281 (10 µM) or ACEA (1 µM), determined by immunostaining (n=4). Each dot represents one mouse and all data are expressed as mean ± s.e.m. Two-sided unpaired Student’s t-test (**B, D** and **F)**, one-way ANOVA followed by Tukey test (**H-I**), two-way ANOVA followed by Tukey test (**E**). Male and female were analyzed independently (**B, D** and **F)**.

### Lack of myeloid CB1 promotes a less pro-inflammatory macrophage phenotype

Our plaque immunostaining analysis indicated a less inflammatory macrophage phenotype in aortic roots of *Apoe^-/-^LysM^Cre^ Cnr1^flox/flox^* mice, which was evident in both males and females (Figure 1G). To follow up on these findings, we performed additional *in vitro* experiments with male BMDMs, considering that the most potent effects of *Cnr1* deficiency were observed in male mice. The transcriptomic profiling of BMDMs by qPCR revealed a downregulation of pro-inflammatory markers (*Irf5, Nos2*) in male *Apoe^-/-^LysM^Cre^ Cnr1^flox/flox^* compared to *Apoe^-/-^LysM^Cre^*BMDMs, while anti-inflammatory markers (*Cx3cr1, Chil3*) were upregulated (Figure 4A). These data were supported by flow cytometry, revealing lower surface expression of pro-inflammatory macrophage activation markers in response to LPS stimulation, namely the receptor CD38 and costimulatory molecule CD80 on *Apoe^-/-^LysM^Cre^Cnr1^flox/flox^* compared to *Apoe^-/-^LysM^Cre^* BMDMs (Figure 4B-C). In agreement with the reduced inflammatory phenotype of *Cnr1-*deficient BMDMs, transcript levels of pro-inflammatory cytokines were less upregulated in response to LPS stimulation (Figure 4D), which was validated at the protein level for the key pro-atherogenic factor IL1-β (Figure 4E). A similar transcriptomic signature was found *in vivo* in peritoneal macrophages of male *Apoe^-/-^LysM^Cre^Cnr1^flox/flox^*compared to *Apoe^-/-^LysM^Cre^* mice collected after 4 weeks WD, whereas only a non-significant difference for reduced *Il1β* expression was detected in females (Supplemental Figure 8A). Moreover, *Apoe^-/-^ LysM^Cre^Cnr1^flox/flox^*BMDMs showed lower oxLDL uptake compared to *Apoe^-/-^LysM^Cre^*BMDMs (Figure 4F). The lower oxLDL uptake was in accordance with a lowered lipid content of plaque macrophages in aortic roots of *Apoe^-/-^LysM^Cre^ Cnr1^flox/flox^*male mice after 4 weeks WD (Supplemental Figure 8B), which supports a role for CB1 in the regulation of macrophage cholesterol uptake or metabolism in atherosclerosis.

**Figure 4.**
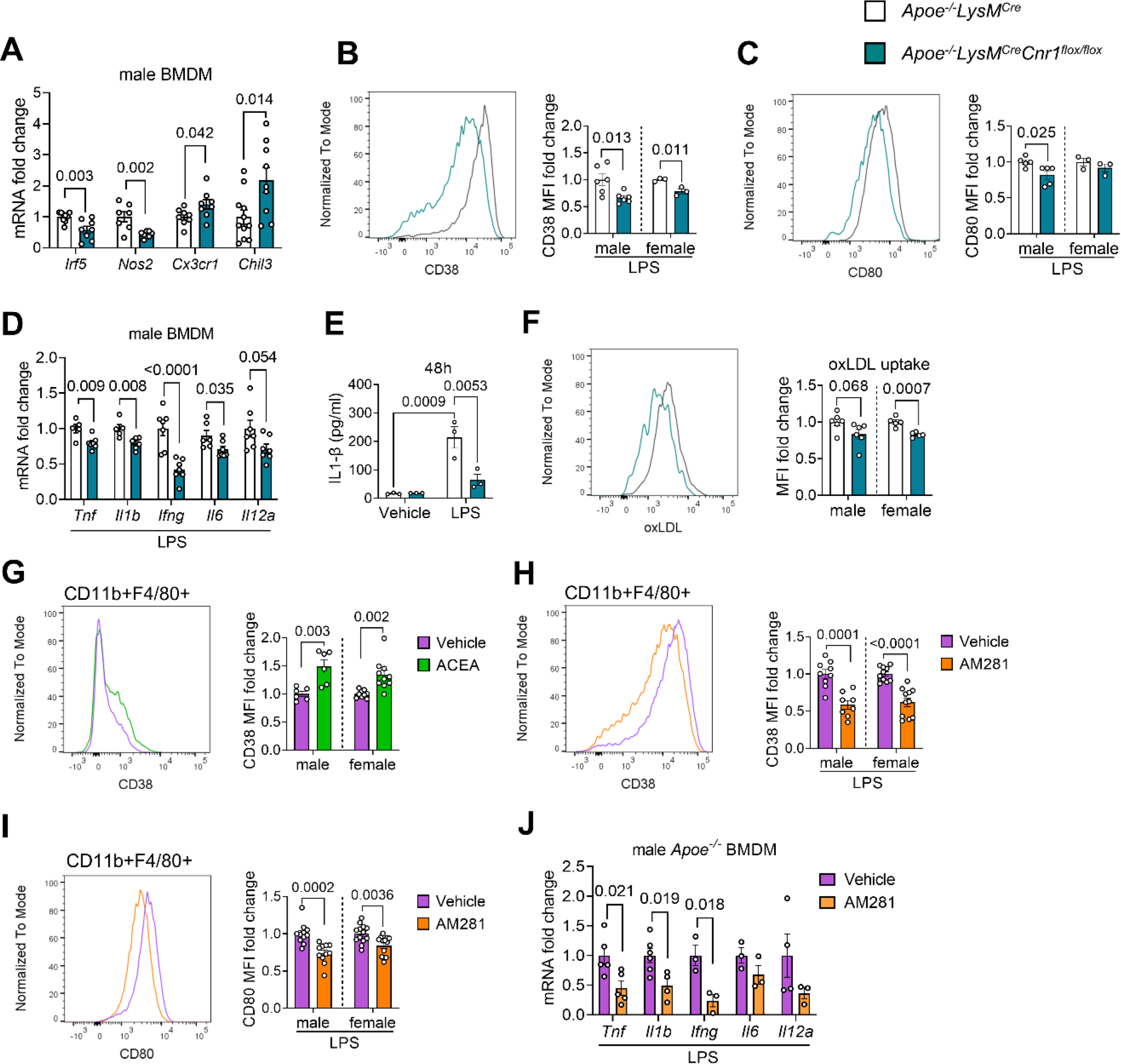
Role of CB1 in inflammatory macrophage polarization and oxLDL uptake. BMDMs were isolated from *Apoe^-/-^LysM^Cre^*and *Apoe^-/-^LysM^Cre^Cnr1^flox/flox^* (**A**-**F**) or *Apoe^-/-^* mice (**G**-**J**). (**A**) Expression levels of macrophage polarization markers in unstimulated BMDMs, determined by qPCR (n=7-12). Flow cytometric analysis of CD38 (**B**) and CD80 (**C**) surface expression of BMDMs after 48 h of LPS (10 ng/mL) stimulation (n=3-6). (**D**) Gene expression levels (qPCR) of pro-inflammatory cytokines in BMDMs after 8 h of LPS-stimulation (n=5–7). (**E**) IL1β secretion of BMDMs after 48 h of LPS-stimulation (n=3). (**F**) Flow cytometric analysis of labelled dil-oxLDL uptake by BMDMs (n=5-6). (**G**) CD38 surface expression on 24 h vehicle- or 1 µM ACEA-treated BMDMs (n=6-10) or (**H**) 24 h LPS-stimulated BMDMs in presence of 10 µM AM281 or vehicle (n=8-12). (**I**) CD80 surface expression on 24 h LPS-stimulated BMDMs in presence of 10 µM AM281 or vehicle (n=8-12). (**J**) Gene expression levels of pro-inflammatory cytokines in *Apoe^-/-^* BMDMs after 8 h LPS stimulation in presence of 10 µM AM281 or vehicle (n=3-5). Each dot represents one biologically independent mouse sample and all data are expressed as mean ± s.e.m. Two-sided unpaired Student’s t-test (**A-D** and **F-J**), two-way ANOVA followed by Tukey test (**E**). Male and female were analyzed independently (**B-C** and **F-I**).

In support of a direct effect of CB1 signaling on macrophage inflammatory reprogramming, stimulating male or female *Apoe^-/-^* BMDMs with the CB1 agonist ACEA upregulated the surface expression of CD38 (Figure 4G), while CB1 antagonism with AM281 inhibited the LPS-induced induction of CD38 and CD80 expression (Figure 4H-I). In addition, the induction of pro-inflammatory cytokine mRNA expression by LPS was suppressed by CB1 antagonism with AM281 (Figure 4J).

### Transcriptomic profiling of BMDMs reveals a profound transcriptional, translational and metabolic regulation by CB1

To gain detailed insights into the molecular pathways elicited by CB1 activation in macrophages, we subsequently performed RNA sequencing of *Apoe^-/-^* BMDMs stimulated with CB1 agonist ACEA. The top gene ontology (GO) biological processes and molecular functions regulated by CB1 were linked to transcriptional regulation, chromatin, and histone modification (Figure 5A-B and Supplemental Figure 9A-C), suggesting that CB1 might control transcriptional regulation through epigenetic modifications. In addition, the transcriptomic signature of ACEA stimulated BMDMs was enriched in pathways linked to myeloid differentiation, leukocyte migration, mRNA processing, actin cytoskeletal organization and spindle organization, which largely fits to our observations in the myeloid *Cnr1* deficiency mouse model. Gene set enrichment analysis (GSEA) further revealed significant associations with inflammatory response, cholesterol transport, and cholesterol efflux (Supplemental Figure 9D).

**Figure 5.**
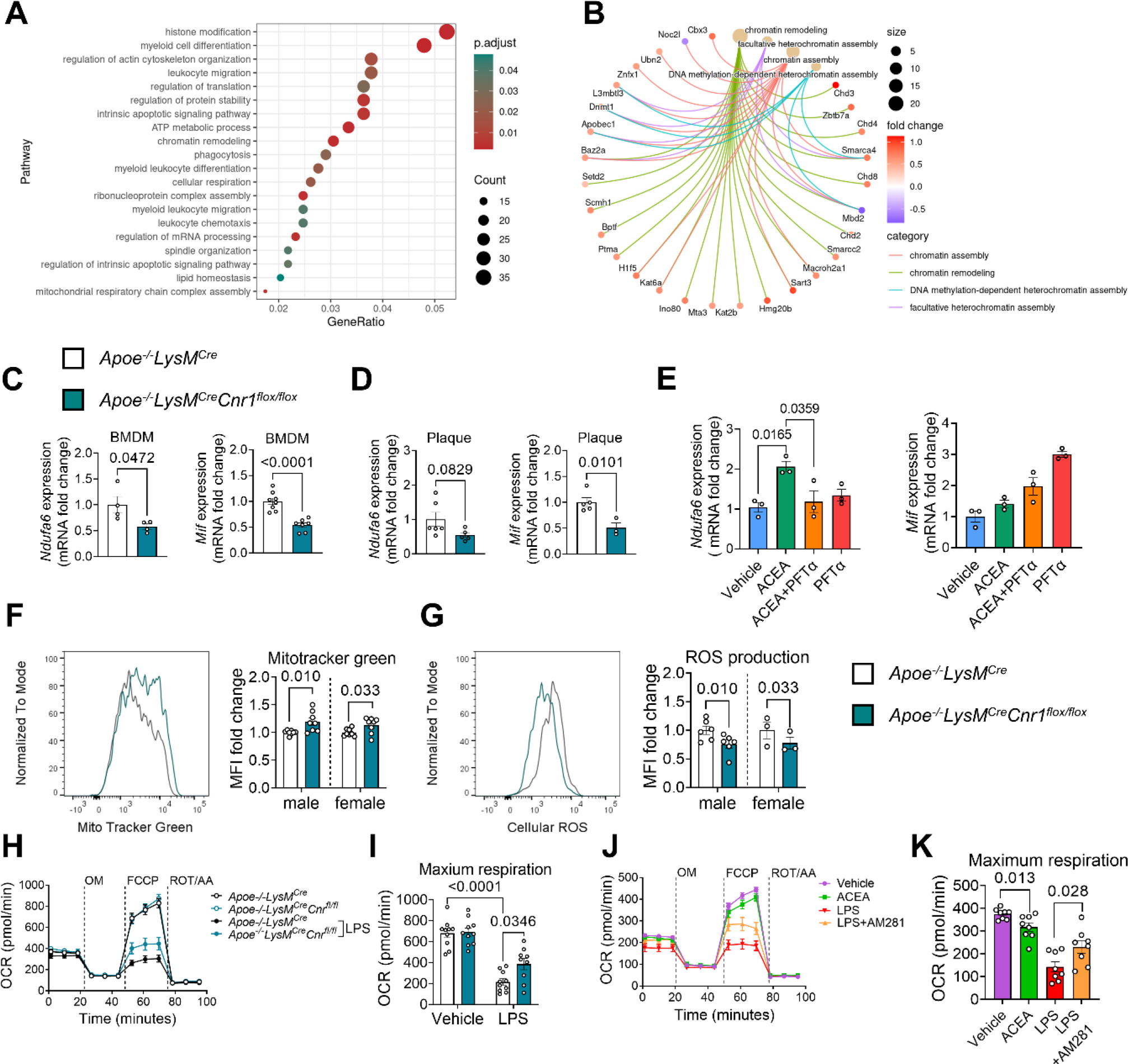
Gene expression profile and mitochondrial effects of CB1 signaling in macrophages. BMDMs were isolated from *Apoe^-/-^* (**A-B, E, J**-**K**) or *Apoe^-/-^LysM^Cre^* and *Apoe^-/-^ LysM^Cre^Cnr1^flox/flox^* mice (**C**-**D, F**-**I**). (**A**) Top regulated biological processes and (**B**) network analysis of chromatin modification pathways and associated DEGs in male BMDMs treated with 1 µM ACEA for 24 h, compared to vehicle. (**C**) Gene expression analysis (qPCR) in BMDMs obtained from baseline male mice (n=4-8) or (**D**) laser capture microdissected plaques of aortic root sections from male mice (n=3-6) subjected to 4 weeks Western diet (WD). (**E**) Gene expression analysis (qPCR) in BMDMs (n=3) treated with 1 µM ACEA for 30 min alone or in presence of p53 inhibitor (PFT-α). (**F**) Flow cytometric analysis of mitochondrial content in BMDMs, determined by MitoTracker™ green staining (n=8), and (**G**) reactive oxygen species (ROS) determined by dihydrorhodamine 123 (DHR123) staining (n=3-6). **(H**–**K)** Mitochondrial respiration in BMDMs (n=8-10 mice), assessed by recording the oxygen consumtion rate (OCR) after sequential injection of oligomycin (OM), carbonyl cyanide 4-(trifluoromethoxy)phenylhydrazone (FCCP), and rotenone (ROT) plus antimycin A (AA). Each dot represents one biologically independent mouse sample and all data are expressed as mean ± s.e.m. wo-sided unpaired Student’s t-test (**C, D, F** and **G**), one-way ANOVA followed by Tukey test (**E** and **K**), two-way ANOVA followed by Tukey test (**I**).

Among the genes upregulated by CB1 stimulation, we found *Mif*, which is an atypical chemokine and well known to play a crucial role in atherosclerosis (Supplemental Figure 9A and C) (21). The *Ndufa6* encoded protein NADH:ubiquinone oxidoreductase subunit A6 is an accessory subunit of the mitochondrial membrane respiratory chain complex I. Accordingly, the GO biological process analysis also indicated a regulation of ATP metabolic process (Figure 5A).

These two key DEGs were confirmed by qPCR, revealing a downregulation of *Ndufa6* and *Mif* in *Apoe^-/-^LysM^Cre^Cnr1^flox/flox^*BMDMs by qPCR (Figure 5C). To further strengthen the *in vivo* relevance of CB1 in regulating the identified key targets, we additionally performed laser capture-microdissection (LCM) of plaques after 4 weeks WD, which revealed a significantly lower *Mif* expression in plaques of *Apoe^-/-^LysM^Cre^Cnr1^flox/flox^* mice, while the reduction in *Ndufa6* levels in LCM plaques did not reach significance (Figure 5D).

In subsequent time course experiments stimulating BMDMs with 1 µM CB1 agonist ACEA, we observed an early upregulation of *Ndufa6* mRNA levels within 30 min, while *Mif* was highly upregulated after 8 h of ACEA stimulation (Supplemental Figure 9E). The top transcription factor binding sites within the *Mif* promoter includes a binding side for p53 (22), which appeared as CB1 target according to the kinase activity profiling (Figure 3G). Pretreatment with the p53 inhibitor PFTα prevented the early upregulation of the *Ndufa6* mRNA level in response to ACEA after 30 min, while the upregulation of *Mif* was not inhibited, but rather enhanced by p53 inhibition (Figure 5E), possibly involving post-transcriptional mechanisms such as mRNA processing and stability.

To follow up on the GO pathway analysis indicating a role for CB1 in mitochondrial oxidative respiration, we addressed a possible regulation of mitochondrial activity by CB1 signaling. Measurement of the mitochondrial content by flow cytometry revealed a significantly higher mitotracker staining intensity in male and female *Apoe^-/-^LysM^Cre^ Cnr1^flox/flox^* compared to *Apoe^- /-^LysM^Cre^* BMDMs (Figure 5F). In support of an improved mitochondrial function, *Cnr1*-deficient BMDMs produced lower levels of reactive oxygen species (Figure 5G). Functional measurement of the oxygen consumption rate (OCR) in BMDMs with the Seahorse extracellular flux analyzer showed no difference in maximum respiration between *Apoe^-/-^ LysM^Cre^ Cnr1^flox/flox^* compared to *Apoe^-/-^LysM^Cre^* BMDM under homeostatic condition; however, *Cnr1* deficiency partially preserved OCR in response to LPS stimulation, resulting in significantly higher maximum respiration rates compared to LPS-treated controls (Figure 5H-I). ACEA treatment of *Apoe^-/-^* BMDMs was sufficient to significantly reduce maximum respiration rates, whereas AM281 partially prevented the LPS-induced decline of OCR (Figure 5J-K).

### Peripheral CB1 antagonism inhibits plaque progression in atherosclerosis

The atheroprotective phenotype of myeloid *Cnr1* deficiency prompted us to address the therapeutic potential of peripheral CB1 antagonism. For this purpose, we decided to use the *Ldlr^-/-^* mouse model of atherosclerosis, which more closely reflects the cholesterol profile in humans compared to the *Apoe^-/-^* model (23). As expected, chronic injection of the peripheral CB1 antagonist JD5037 for 4 weeks improved some metabolic parameters, including lower plasma cholesterol levels in both males and females, lower plasma triglyceride levels in males and significantly reduced body weight increase only evident in males (Supplemental Figure 10A-D).

Moreover, the 4-week treatment was sufficient to inhibit early-stage plaque progression in aortic roots and arches of male mice, whereas female mice treated with either JD5037 or vehicle had comparable plaque sizes (Figure 6A-F). The reduced plaque growth in male aortic roots was related to a lower macrophage accumulation, proliferation, and inflammatory polarization as indicated by a lower number of iNOS+CD68+ plaque macrophages (Figure 6G-I and Supplemental Figure 10E). A lower surface CD80 expression level on peritoneal macrophages was found in both male and female mice receiving JD5037, indicative of a decreased inflammatory phenotype (Supplemental Figure 10F). The differential effects of peripheral CB1 antagonism on female peritoneal versus plaque macrophages might be related to higher local concentrations of JD5037 at the peritoneal injection site as well as distinct transcriptomic signatures of macrophages depending on their tissue origin (24).

**Figure 6.**
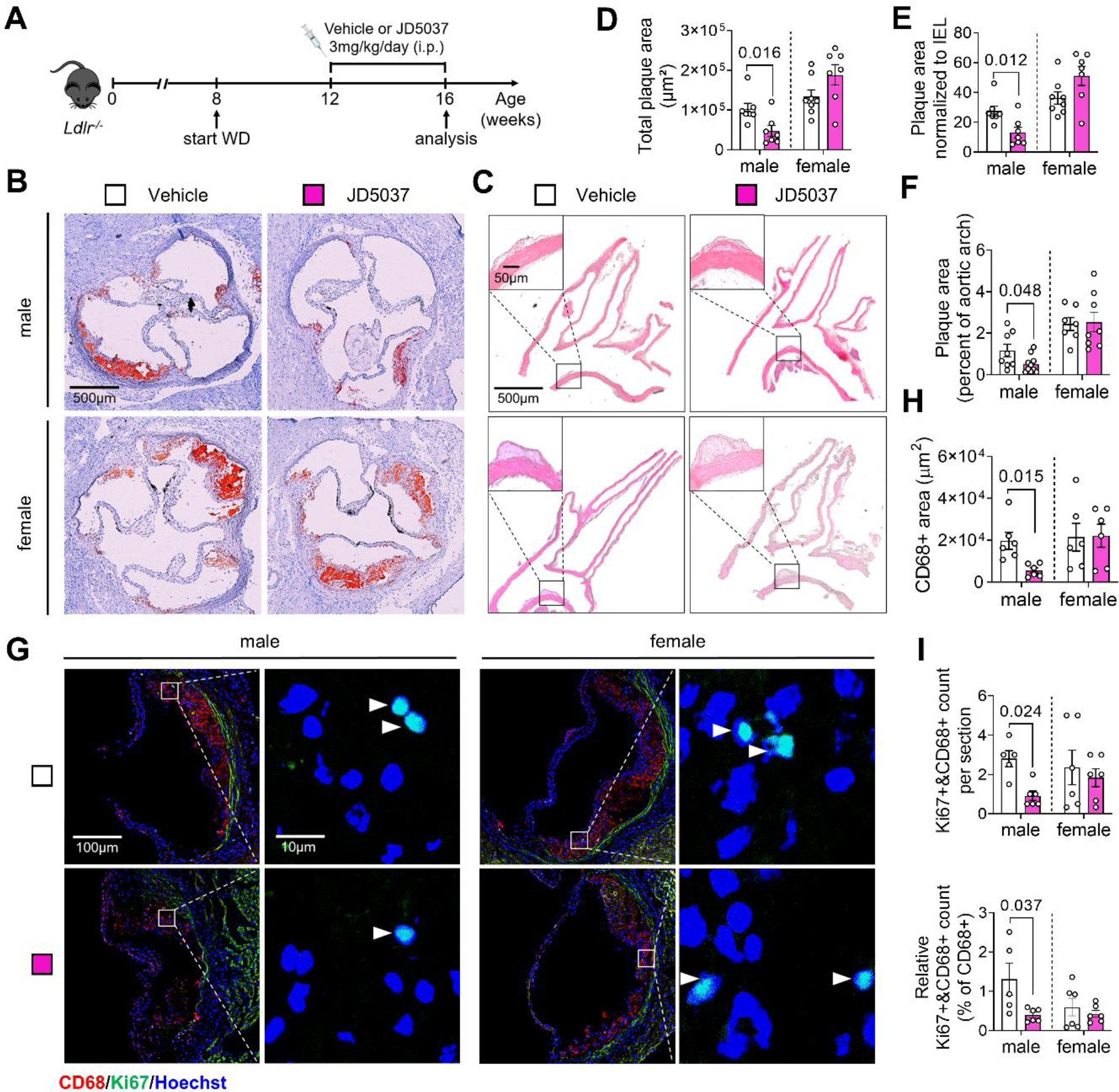
Effect of chronic peripheral CB1 antagonist administration on atherosclerosis and macrophage proliferation. (**A**) Experimental scheme. *Ldlr*^−/−^ mice were fed with Western diet (WD) for 8 weeks and subjected to daily i.v. injections of JD5037 (3 mg/kg) or vehicle for the last 4 weeks. (**B**) Representative images of aortic root cross-sections stained with ORO. (**C**) Representative images of HE-stained aortic arch longitudinal sections. (**D**) Quantification of absolute lesion area within aortic roots (n=7-8 mice) or (**E**) normalized to IEL (n=7-8). (**F**) Quantification of the plaque area in aortic arch sections (n=7-8). (**G**) Representative images of proliferating (Ki67+, white arrowheads) macrophages (CD68+) in aortic root plaques; nuclei were counterstained with Hoechst. Scale bar: 100 μm (left) and 10 µm (right). (**H**) Quantification of lesional macrophages (n=6) and **(I)** total as well as relative counts of proliferating plaque macrophages (n=5-6). Each dot represents one biologically independent mouse sample and all data are expressed as mean ± s.e.m. Two-sided unpaired Student’s t-test (**D-F, H-I**). Male and female were analyzed independently.

### Expression of CNR1 in advanced human atherosclerosis

Finally, to address a possible implication of CB1 signaling in the regulation of inflammatory gene expression, mitochondrial function, and proliferation in human atherosclerotic plaques, bulk RNA seq transcriptomic data from patients undergoing carotid endarterectomy (CEA) were analyzed (25). *CNR1* transcript levels inversely correlated with the proliferation marker *MKI67* and several genes regulated in a CB1-dependent manner in murine BMDMs, including the mitochondrial membrane respiratory chain *NDUFA6* as well as the inflammatory genes *CCR5, MIF, CD74, IL1B* and *IL6* (Figure 7). This may support a direct relationship between CB1 and key cellular pathways in human plaque pathophysiology as well as mechanisms involved in lesion destabilization.

**Figure 7.**
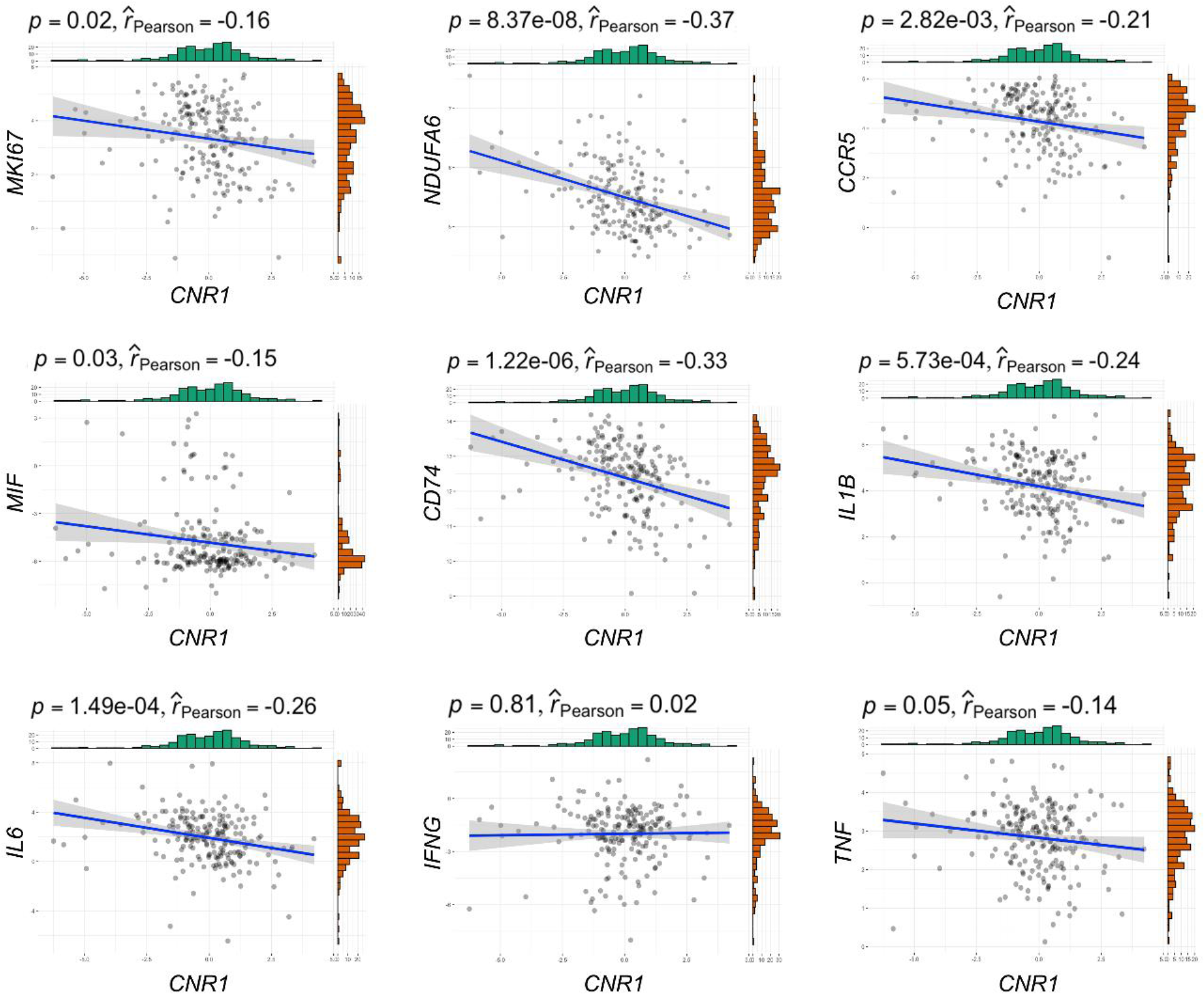
Correlation analysis between *CNR1* and key markers of proliferation, mitochondrial oxidative metabolism and inflammation in human plaques. Bulk RNAseq data from human carotid plaques were used for linear regression analysis, based on normalized gene expression levels of selected genes. Scatter plots and regression lines are shown (n = 203 samples). Pearson correlation coefficients (r) and P values are presented in the graphs.

## Discussion

Using a targeted genetic depletion strategy and a pharmacological blocking approach, we here provide previously unknown insights into the molecular mechanisms of myeloid cell CB1 signaling in the context of atherosclerosis. We found pleiotropic effects of CB1 signaling in circulating monocytes, plaque macrophages, peritoneal macrophages, and *in vitro* cultured BMDMs. The biological processes regulated by CB1 in monocytes and macrophages include recruitment and chemotaxis, proliferation, inflammatory cytokine production, oxLDL uptake, and mitochondrial oxidative respiration. The underlying molecular regulation appears to be multifold, including p53-dependent transcriptional regulation and possibly chromatin accessibility as well as post-transcriptional mechanisms. Remarkably, the atheroprotective effects of myeloid CB1 deficiency were more pronounced in male mice, at least in early disease stage. At advanced stage, a significantly smaller plaque was also found in female mice, albeit this difference was only detectable in the aortic arch. This indicates that the impact of *Cnr1* deficiency on the atherosclerotic phenotype differs depending on the disease stage, the specific location within the arterial tree and sex. The underlying reason is likely that different cellular key players and biological processes contribute to the various disease stages, which are differentially affected by sex hormones and other sex-specific determinants. During early atherogenesis, monocyte recruitment, macrophage differentiation into foam cells, and local proliferation are key processes driving plaque formation. At a more advanced stage, macrophage apoptosis, clearance of apoptotic cell debris and necroptosis contribute to plaque progression (16, 26). Interestingly, the proliferation rate *per se* was lower in female compared to male macrophages, which appears to be attributable to an inhibitory effect of the female sex hormone estradiol on cell proliferation, as previously reported, among other in monocytic leukemia cells and vascular smooth muscle cells (27, 28).

The effects of CB1 signaling on cellular proliferation might be related to a regulation of cellular cholesterol metabolism, as supported by our GSEA data, given that genetic mutations in cholesterol uptake receptors or efflux critically affect myeloid cell proliferation (29, 30). In line with this, CRISPR/Cas screens in human cell lines, accessible through the bioGRID repository for interaction datasets (31), support a link between *Cnr1* and cell proliferation (https://orcs.thebiogrid.org/Gene/12801). In a previous *in vitro* study, synthetic cannabinoids reduced the expression of cholesterol efflux transporter ABCA1 in RAW264.7 cells, while increasing scavenger receptor CD36 expression, which mediates uptake of modified LDL. The effect was sensitive to pretreatment with CB1 antagonist/inverse agonist AM251 (14). Human genetic studies have identified a causal variant in the *CNR1* promoter that is linked to high-density lipoprotein cholesterol levels (32), which supports a rather complex role for CB1 signaling in the regulation of cholesterol metabolism. In support of a regulation of macrophage cholesterol metabolism by CB1 in atherosclerosis, we found reduced oxLDL uptake by BMDMs and a lower amount of lipid-laden plaque macrophages in aortic roots of *Apoe^-/-^LysM^Cre^ Cnr1^flox/flox^* male mice after 4 weeks WD.

Furthermore, BMDMs lacking CB1 produced lower amounts of IL1β, which indicates a reduced inflammasome activation in plaques of *Cnr1*-deficient mice. This is in accordance with a previous study linking CB1 signaling to inflammasome activation in pancreatic macrophages (13). The reduced cholesterol accumulation in plaque macrophages might contribute to the less inflammatory and more stable plaque phenotype, given that excessive cholesterol accumulation impairs the capacity of macrophages to clear apoptotic cells. This is also supported by our GSEA analysis, revealing that CB1 activation in macrophages is linked to an upregulation of genes linked to inflammatory response and cholesterol. Impaired clearance of apoptotic cells by plaque macrophages, leading to secondary necrosis, and extracellular lipid accumulation contribute to the formation of a necrotic core. Secondary necrosis as well as alternative, non-apoptotic cell death pathways (such as necroptosis or pyroptosis) occurring within advanced lesions further entrain the local inflammatory milieu, thereby contributing to plaque destabilization (33). Hence, the reduced inflammatory phenotype of *Cnr1*-deficient macrophages is in agreement with a smaller necrotic plaque size in *Apoe^-/-^LysM^Cre^ Cnr1^flox/flox^* mice.

The kinase profiling array revealed a regulation of p53 activity and several cyclin-dependent kinases in response to CB1 stimulation, which are well known to be involved in cell cycle regulation (34). A link between CB1 and p53 activity has been previously described in an experimental study of diet-induced hepatic steatosis (35). The p53-dependent transcriptional regulation might be a central mechanism in our atherogenic mouse model with myeloid *Cnr1* deficiency, in addition to epigenetic mechanisms. In fact, p53 has been previously linked to macrophage recruitment, proliferation, and smaller necrotic core size (36–38). Furthermore, latest findings have unveiled that *TP53*-mediated clonal hematopoiesis confers increased risk for incident atherosclerotic disease (39).

The chemokine receptor analysis at protein and mRNA level provided strong evidence for a direct regulation of CCR1 and CCR5 expression by CB1. The CB1-dependent regulation of *Ccr5* was further confirmed by bulk RNA sequencing of BMDMs stimulated with CB1 agonist. Among the top DEGs regulated by CB1, we found *Mif* and *Cd74* as one of its cognate receptors. The *Mif* gene encodes for macrophage migration-inhibitory factor (MIF), which is an inflammatory cytokine and atypical chemokine lacking the typical chemokine-fold and conserved cysteines (21). MIF is upregulated in atherosclerosis, promoting atherogenic leukocyte recruitment and vascular inflammation through the chemokine receptors CXCR2, CXCR4, CD74 and ACKR3 (21). CD74, also known as MHC class II invariant chain, has been involved in diverse immunological functions such as cancer cell proliferation, by serving as high affinity MIF receptor (40).

The GO analysis further revealed a regulation of mitochondrial ATP production, which was supported by functional metabolic measurements. It is well documented that pro-inflammatory macrophage polarization leads to suppression of oxidative respiration (41), which was partially blocked by CB1 antagonism. This suggests that LPS triggers an acute release of endocannabinoids by the macrophages, which activate their own CB1 receptors. The inhibitory effect of the synthetic CB1 agonist ACEA on the macrophage mitochondrial respiration was less pronounced compared to the LPS effect, which may indicate that CB1 signaling alone is insufficient for metabolic reprograming of macrophages. Another explanation might be that the exogenous agonist is less efficient to affect mitochondrial metabolism compared to endogenously produced ligands in response to LPS stimulation. In this context, it is also noteworthy that mitochondrial CB1 receptors have been identified in the brain and are linked to neuronal energy metabolism (42). However, given that CB1 detection at the protein level is hampered by the lack of specific antibodies, it remains a question to be solved in future studies if mitochondrial CB1 signaling may possibly play a role in the regulation of macrophage metabolism.

Peripheral CB1 antagonism had multiple anti-inflammatory and metabolic effects in our experimental atherosclerosis mouse model, and it is likely that the anti-atherogenic effects are more complex and not solely due to effects on myeloid cells. The protective actions might be partially related to cholesterol lowering and possible effects on other cellular key players involved in the pathophysiology of atherosclerosis, which deserves further investigation but is beyond the scope of the present study. Nevertheless, the rationale of this experiment was to clarify whether key cellular effects observed in mice with myeloid *Cnr1* deficiency can be modulated by chronic treatment with a peripheral CB1 antagonist. Indeed, we were able to reproduce inhibitory effects on plaque macrophage accumulation, proliferation, and inflammatory polarization by chronic treatment with a peripheral CB1 antagonist.

The regression analysis of transcriptomic plaque data from patients undergoing CEA revealed an inverse correlation between *CNR1* expression and markers of proliferation, oxidative phosphorylation, and several inflammatory genes. These associations may support a direct relationship between CB1 and key cellular pathways in human plaque pathophysiology. Based on our findings in the mouse model and *in vitro* experiments in BMDMs, the inverted correlations are in part opposite to the anticipated CB1-dependent stimulation of macrophage proliferation and inflammatory phenotype. However, the *CNR1* transcript levels were determined via bulk RNAseq analysis of entire human plaques and therefore may not only reflect the *CNR1* expression in macrophages, given that other cell types, such as endothelial cells and smooth muscle cells also contribute to the total plaque *CNR1* expression levels (43, 44). In addition, it is possible that chronic CB1 stimulation in inflammatory plaques limits expression, possibly due to receptor desensitization, as previously described in mice with chronically elevated endocannabinoid levels due to pharmacological inhibition of the metabolizing enzyme monoacylglycerol lipase (45). Furthermore, it is noteworthy that a human genome-wide association study identified a single nucleotide polymorphism (*rs75205693*) in the *CNR1* gene, which is associated with DNA methylation (46).

A limitation of our study is that we only focused on the molecular effects of CB1 in monocytes and macrophages in our study. However, we cannot exclude that lack of CB1 signaling in neutrophils may partially contribute to the atheroprotective phenotype in our mouse model with myeloid *Cnr1* depletion. We did not observe changes in CXCR2 surface expression levels on circulating neutrophils of mice lacking myeloid CB1. In addition, the immunostaining analysis of neutrophils within the plaques after 4 weeks WD did not reveal any difference between the genotypes. Furthermore, the observed sex-specific differences in macrophage phenotypes, both in experimental atherosclerosis *in vivo* and in BMDMs *in vitro*, deserve a more depth investigation in follow up studies. Elucidating the underlying mechanisms, which are likely more complex than only attributable to female sex hormones, should be addressed in suitable experimental models such as the “four core genotypes” (FCG) mouse model (47). Likewise, investigating sex-specific dimorphism in clinical studies certainly deserves more attention in the future.

In conclusion, our data suggest that inhibiting CB1 signaling in monocytes and macrophages exhibits multiple atheroprotective effects: blocking CB1 inhibits the recruitment of monocytes to atherosclerotic plaques and dampens the local proliferation of plaque macrophages, and it reduces inflammatory cytokine release, oxLDL uptake, and improves mitochondrial oxidative respiration. The biological function of CB1 signaling in macrophages seems to be at least in part sex-dependent, playing a more pronounced pro-atherosclerotic role of myeloid CB1 in male, which appears to be less relevant in female mice. It further highlights the need to consider the biological sex as an important variable in preclinical studies. The clinical relevance for this possible sex-specific effect of CB1 signaling in monocytes and macrophages deserves further investigation.

## Methods

### Animal model of atherosclerosis

Mice harboring loxP-flanked *Cnr1* alleles (*Cnr1^flox/flox^*) (48) were crossed with apolipoprotein E– deficient mice (*Apoe^-/-^* mice, strain #002052, The Jackson Laboratory) expressing Cre recombinase under the control of the myeloid cell-specific lysozyme M promoter (*LysM^Cre^*) to obtain mice with a myeloid cell-specific deletion of *Cnr1*. Male and female *Apoe^-/-^LysM^Cre(+/-)^, Apoe^-/-^Cnr1^flox/flox^ and Apoe^-/-^LysM^Cre(+/-)^Cnr1^flox/flox^* mice aged 8 weeks were fed with Western diet (WD; 0.2% cholesterol, Ssniff, TD88137) for either 4 or 16 weeks. In other experiments, *Ldlr-/-* mice were fed with WD for 4 weeks and subsequently randomly assigned into 2 groups receiving daily intraperitoneal injections of either JD5037 (3 mg/kg) or vehicle (10% DMSO, 40% PEG3000, 5% Tween-80 and 45% saline) for 4 weeks with continued WD feeding. Organs for baseline phenotyping were collected from 8 weeks old mice kept on normal chow diet. Animals were housed in individual ventilated cages in groups of 4 to 6 mice in 12-hour light– dark-cycle, air-conditioned (23°C and 60% relative humidity) with free ad libitum access to food pellets and tap water. Mice were euthanized under deep anesthesia with ketamine/xylazine, in order to collect blood via cardiac puncture and perfusion of organs before harvesting the heart, aorta, spleen, femurs, and livers.

### Plasma cholesterol and triglyceride measurement

Total murine plasma cholesterol and triglyceride concentrations were measured with colorimetric assays (CHOD-PAP and CPO-PAP, Roche) using a microplate reader (Infinite F200 PRO, Tecan).

### Plaque analysis in descending aorta

Oil-Red-O (ORO) stainings of the aorta including abdominal and thoracic aorta were performed after en face preparation and fixation in 4% PFA. The tissue was stained for 30 min in ORO solution (0.5% in isopropanol, Sigma Aldrich/Merck) at RT. Next, aortas were washed in 60% isopropanol until unspecific ORO-derived staining was removed from the endothelium. The aortas were rinsed briefly with running tap water before mounting in Kaiser’s glycerin gelatine mounting media. Tilescan images were acquired with a DM6000B fluorescence microscope. The lesions were quantified via Leica Application Suite LAS V4.3 software and calculated as percentage of total aortic surface.

### Histology and immunofluorescence

Hearts were embedded in Tissue-Tek O.C.T. compound (Sakura) for cutting 5 µm cross-sections at the level of the aortic roots. Aortic arches were fixed in 4% PFA and embedded in paraffin for longitudinal sectioning (5 µm). Aortic root sections were stained with ORO for lesion size quantification. Per mouse, 8 sections with a distance of 50 µm from each other were analyzed, and mean values per heart calculated. Aortic arch slides were deparaffinized and stained with hematoxylin and eosin for lesion size quantification relative to the total inner vessel area. Plaque collagen content was assessed by Masson’s trichrome staining (stain kit Sigma HT15) of deparaffinized aortic arch sections. Per arch, 4 sections at 50 µm distance from each other were analyzed. Digital images were acquired with a brightfield microscope (DM6000B) connected to a digital camera (DMC6200, Leica). The lesion size was quantified using Leica Application Suite LAS V4.3 software.

Immunostainings were performed on 5 µm thick cryosections of the aortic root. After fixation with pre-cooled acetone and blocking nonspecific binding (1% BSA in PBS), sections were incubated with antibodies against CD68, iNOS, Ki67 and Ly6G (Supplementary Table 1-2). Negative staining controls were performed with corresponding isotypes. The primary antibodies were detected with anti-rat Cy3 and anti-rabbit AF488 conjugated secondary antibodies (Supplementary Table 3). For staining of lipid droplets within macrophages, CD68 staining was followed by Nile Red (N3013, Sigma-Aldrich) staining for 5 minutes. Nuclei were counterstained with Hoechst 33342. Images were taken with a fluorescence microscope (DM6000B) connected to a monochrome digital camera (DFC365FX, Leica) equipped with Thunder technology for computational clearing, and analyzed with the Leica Application Suite LAS V4.3 software or ImageJ software. Per heart, 3 aortic root sections were analyzed, and mean values per section calculated.

### *In situ* hybridization

To determine the presence of CB1 in macrophage, fluorescent in situ hybridization was performed by using a murine *Cnr1* probe (VB6-17606, Affymetrix). Aortic root cryosections were collected on RNAse free slides (pretreated with RNAse ZAP). RNase-free requirements were maintained up to post-hybridization steps. The sections were prefixed in 4% PFA for 5 min at RT and washed 3 times with PBS for 5 min. The tissue sections were treated with pre-warmed 10 μg/ml proteinase K (diluted in PBS) for 5 min at RT, followed by post-fixing with 100% ethanol for 1 min. After washing 3 times for 10 min with PBS, the sections were transferred to an RNAse free chamber. The ViewRNA™ Cell Plus Assay-Kit (Invitrogen) was used for target probe hybridization by incubating the sections at 40° C for 2 h. The hybridization probe was mixed with probe set diluent to a final concentration of 5 μg/ml. Following the hybridization step, the sections were washed 3 times with wash buffer at RT for 5 min. Signal amplification was performed by incubating with the pre-amplifier mix for 30 min at 40° C, washing and incubating with the amplifier mix for 1 h at 40°C. After washing, the sections were incubated for 1 h at 40°C with the working label probe mix. Following the in-situ hybridization, the sections were washed, incubated with blocking buffer and stained with CD68 antibody and Hoechst as described above. Images were taken using a Leica DM 6000B microscope.

### Laser capture microdissection

Aortic root sections (8-10 sections per mouse, 10 µm thick) from *Apoe^-/-^LysM^Cre^* and *Apoe^-/-^ LysM^Cre^Cnr1^flox/flox^* male mice collected after 4 weeks WD were mounted on RNase-free membrane metal frame slides (POL FrameSlide, Leica Microsystems). Following deparaffinization under RNase-free conditions, sections were dried. Lesions were cut using a laser microdissection system (CTR6000, Leica Microsystems) attached to a microscope (LMD7000, Leica Microsystems) and collected in lysis buffer of the RNA isolation kit (PAXgene Tissue miRNA kit, PreAnalytix).

### Monocyte adoptive transfer

Monocytes were isolated from the bone marrow of *Apoe^-/-^LysM^Cre^* and *Apoe^-/-^LysM^Cre^Cnr1^flox/flox^* mice by magnetic separation monocyte Isolation Kit (130-100-629, Miltenyi Biotec). The purified monocytes were labelled with carboxyfluorescein diacetate succinimidyl ester (CFSE, 10 μM, V12883, ThermoFisher) and 2.5×10^6^ cells injected intraveneously via the tail vein. Both donor mice and *Apoe^-/-^* recipient mice were subjected to 4 weeks WD before adoptive cell transfer. After 48h, the aorta and hearts were collected after organ perfusion to detect recruited CFSE+ monocytes within aortic root cryosections by fluorescence microscopy (DM6000B) and digested aortas using flow cytometry.

### Flow cytometry

Femurs were centrifuged at 10,000 × g for 1 min after exposure of the distal metaphysis to collect the bone marrow cells. Bone marrow and blood erythrocytes were lysed with ammonium-chloride-potassium (ACK, NH_4_Cl (8,024 mg/l), KHCO_3_ (1,001 mg/l) EDTA.Na_2_·2H_2_O (3.722 mg/l) buffer for 10 min at RT. The cell suspensions were subsequently blocked for 5 min with Fc-CD16/CD32 antibody and then stained for 30 min in the dark at 4° C with antibodies (Supplementary Table 4) to identify myeloid and lymphoid cell subsets. After gating for singlets and CD45+, the CD11b+ myeloid subsets were further gated as following: CD115+Ly6G-(monocytes), CD115-Ly6G+ (neutrophils). Surface expression levels of chemokine receptors CCR1, CCR5 and CXR2 on myeloid cell subsets were expressed as geometric mean fluorescence intensity (MFI). For measuring proliferation, cells were permeabilized (eBioscience, 00-8333-56) prior to staining with anti-Ki67. Flow cytometry data were acquired on a BD FACSCantoII flow cytometer (BD Biosciences) and analyzed with FlowJo v10.2 software (Tree Star, Inc). Total cell counts in blood and bone marrow samples were determined using CountBright absolute counting beads.

### Flow cytometric cell sorting

Blood samples from *Apoe^-/-^* mice were subjected to red blood cell lysis and antibody stainings as described above for sorting CD19+ B cells, CD3+ T cells, CD115+ monocytes, and Ly6G+ neutrophils using a BD FACSAria III Cell Sorter (BD Biosciences). The sorted cells were deep frozen in 2x TCL buffer plus (Qiagen) plus 1% beta-mercaptoethanol.

### *In vitro* assays with bone marrow-derived macrophages

Bone marrow cells were collected from tibia and femur by centrifugation (8600x g for 1min at 4 °C) and then subjected to red blood cell lysis using ACK buffer. After washing, the cells were resuspended in RPMI-1640 (Gibco Life Technologies) medium supplemented with 10% FCS, 2 mM L-glutamine, penicillin (100 U/mL), streptomycin (100 μg/mL) and 10 % filtered L-929 cell-conditioned medium as a source of M-CSF. The cells were plated into 12-well-plates, and the medium was changed every 2–3 days for a total of 7-10 days to allow differentiation into bone marrow-derived macrophages (BMDMs). For immunostainings, cells were plated on sterile cover slips in 24-well plates. In some experiments, BMDMs were treated with estradiol (10 µM), LPS (10 ng/mL), CB1 agonist ACEA (1 µM), or CB1 antagonist AM281 (10 µM). In case of combined LPS treatment with CB1 blocking, AM281 was added 15 min prior to LPS. The p53 inhibitor pifithrin-α (PFT-α; 25 µM) was added 5 min before stimulating the cells with ACEA or AM281.

For ROS detection, BMDM were incubated with DHR123 (2.5 µg/ml; 851000, Cayman) in RPMI 1640 (plus Glutamine) containing 0.1% BSA and 0.1 mM HEPES for 20 min at 37°C. Then, the samples were immediately acquired with a FACSCantoII to detect cellular ROS levels as green fluorescence intensities. For mitochondrial staining, BMDMs were incubated with prewarmed (37°C) staining solution containing MitoTracker (ThermoFisher) at a concentration of 50 nM for 30 min and subsequently measured with a FACS Canto-II. For assessing oxLDL uptake, BMDMs were incubated with 1 µg/ml Dil-oxLDL in complete RPMI medium for 30min. After loading, cells were washed twice with prewarmed PBS before detaching with prewarmed citrate buffer (135mM KCl, 15 mM sodium cirate). The amount of intracellular Dil-oxLDL was measured with a FACS Canto-II. Cell proliferation rates were measured by intracellular Ki67 staining and flow cytometry as described above. IL1β levels in BMDM supernatants were measured by DuoSet ELISA (R&D Systems) with a Tecan Infinite F200 PRO microplate reader. For detection of nuclear p53, cells grown on cover slips and treated for 1h with ACEA, AM281 or vehicle were fixed with acetone and subsequently immunostained with primary antibody against phospho-p53 and subsequently with anti-rabbit-AF488. Images were taken with a fluorescence microscope (DM6000B) connected to a monochrome digital camera (DFC365FX, Leica) equipped with Thunder technology for computational clearing, and analyzed with the Leica Application Suite LAS V4.3 software or ImageJ software. Throughout all experiments, each independent biological replicate was generated with separate BMDM donor mice.

### Quantitative real time PCR

Total RNA from BMDM was isolated (peqGOLD, 13-6834-02, VWR Life Science) and reverse transcribed (PrimeScript RT reagent kit, TaKaRa). Real-time qPCR was performed with the QuantStudio™ 6 Pro Real-Time PCR System (ThermoFisher) using the GoTaq Probe qPCR Master Mix (Promega). Primers and probes were purchased from Life Technologies (Supplemental Table 5). Target gene expression was normalized to hypoxanthine guanine phosphoribosyltransferase (*Hprt*) gene expression and represented as fold change relative to the control group.

### Droplet digital PCR

PCR was performed on QX200 Droplet Digital PCR (ddPCR™) system from Bio-Rad. The reaction mixture was prepared by combining 2× ddPCR Mastermix (Bio-Rad), 20× primer and probes (final concentrations of 900 nM and 250 nM, respectively; Integrated DNA Technologies) and cDNA template in a final volume of 20 μl (Supplementary Table 5). Then, each ddPCR reaction mixture was loaded into the sample well of an eight-channel disposable droplet generator cartridge (Bio-Rad). A volume of 60 μl of droplet generation oil (Bio-Rad) was loaded into the oil well for each channel. The cartridge was placed into the droplet generator (Bio-Rad), and the droplets were collected and transferred to a 96-well PCR plate. The plate was heat-sealed with a foil seal, placed on the thermal cycler and amplified to the endpoint (40 cycles). Subsequently, the 96-well PCR plate was loaded on the droplet reader (Bio-Rad) and the analysis was performed with QuantaSoft analysis software (Bio-Rad).

### RNA sequencing

BMDMs were isolated (n=5 donor mice) and cultured as described above and stimulated with ACEA for 24 hours. Total RNA was isolated from BMDMs using the RNeasy Mini kit with an on-column DNase I treatment according to the manufacturer’s instructions (Qiagen). RNA concentration was determined using a Qubit fluorometer (Qiagen) while RNA quality was evaluated using an Agilent Biolanalyzer 2100 system (Agilent Technologies, CA, USA) and subsequently used to construct strand-specific libraries using the KAPA mRNA HyperPrep kit according to the manufacturer’s instructions. Briefly, mRNA was captured, fragmented and primed for cDNA synthesis followed by adapter ligation for identification. Libraries were pooled and diluted to 30 nM prior to sequencing on an Illumina HiSeq 4000 instrument (Illumina) at a depth of 20 million single ended 50 base pair reads. Reads were aligned to the mouse genome mm10 using STAR 2.5.2b with default settings. BAM files were indexed and filtered on MAPQ > 15 with SAMtools 1.3.1. Raw tag counts and RPKM (reads per kilo base per million mapped reads) values per gene were summed using HOMER2’s analyzeRepeats.pl script with default settings and the -noadj or -rpkm options for raw counts and RPKM reporting, respectively. Datasets were filtered to select genes expressed in at least 10% of samples. Differential gene expression analysis was calculated by using DESeq2 (Version 1.37.4) Bioconductor package in R version 4.2.0 (2022-04-22) under Ubuntu 20.04.3 LTS system, based on the negative binomial distribution (49). Size factors and dispersion of samples were estimated first, then Negative Binomial GLM fitting and Wald statistics calculation was performed by DESeq2. Differential expression genes were identified by criteria: adjusted P value < 0.10 and fold changes > 1.5, based on the calculation results of DESeq2. Gene ontology enrichment analysis was performed using the enrichplot (Version 1.17.2) Bioconductor package (50) with differential expression genes (only filtered by adjusted P value < 0.10). Heatmap was drawn by pheatmap package (Version 1.0.12). Volcano plot was drawn by ggplot2 package (Version 3.4.0). Cneplots and dotplots were drawn by enrichplot packge (Version 1.17.2). The Gene set enrichment analysis (GSEA) was done on GSEA software v4.3.2 for Windows with M2 curated gene sets and M5 ontology gene sets from mouse collections and C2 gene sets from human collections (51, 52). The permutation type was gene_set. The significant pathways were chosen by following criteria: normalized enrichment score (NES) < -1 or > 1, false discovery rate (FDR) < 25% and nominal P value < 0.05.

### Phospho kinase array

BMDMs isolated from 4 donor mice were treated with ACEA for 60 min and then washed once in ice-cold PBS, and lysed for 15 min on ice using M-PER Mammalian Extraction Buffer containing Halt Phosphatase Inhibitor and EDTA-free Halt Protease Inhibitor Cocktail (1:100 each; Thermo Scientific). Lysates were centrifuged for 15 min at 16,000 × g at 4°C. Protein quantification was performed with Pierce™ Coomassie Plus (Bradford) Assay according to the manufacturer’s instructions. Serine-Threonine kinase (STK) profiles were determined using the PamChip® peptide Ser/Thr Kinase assay microarray systems on PamStation®12, respectively (PamGene International). Each STK-PamChip® array contains 144 individual phospho-site(s). Per array, 1.0 μg of protein and 400 μM ATP were applied together with an antibody mix to detect the phosphorylated Ser/Thr. The spot intensity was quantified (and corrected for local background) using the BioNavigator software version 6.3 (PamGene International). Upstream Kinase Analysis, a functional scoring method (PamGene), was used to rank kinases based on combined specificity scores (based on peptides linked to a kinase, derived from six databases) and sensitivity scores (based on treatment-control differences). The heatmap contains the kinases with a median final scores >1.2 and p-values were adjusted for multiple comparisons by the false discovery rate (FDR). Pathway enrichment analysis was implemented on the differentially expressed kinases (median final scores >1.2 and adjusted p-value <0.05) with the use of the R packages ReactomePA (50).The biological complexities in which these kinases correlate with multiple annotation categories were visualized in a network plot with the help of the R package Enrichplot Version 1.4.0. 2019 (10.18129/B9.bioc.enrichplot).

### Extracellular flux analysis

Mitochondrial respiration was assessed using the Seahorse Cell Mito Stress Test in a Seahorse Analyzer XFe24 (Agilent) with some adjustments to the manufacturer’s protocol. Briefly, 4×10^5^ BMDM cells were transferred to a 24-well Seahorse plate after 7 days of differentiation. BMDM were incubated with CB1 agonist ACEA (1 µM), CB1 antagonist AM281 (10 µM), LPS (10 ng/ml), or vehicle for 48h. Prior to the assay, the culture medium was replaced with Seahorse medium (XF RPMI medium, pH 7.4, supplemented with 10 mM glucose, 1 mM pyruvate and 2 mM L-glutamine) and the cells were incubated for 60 min at 37 °C without CO_2_. The oxygen consumption rate (OCR) was measured (3 min mix, 2 min wait and 3 min measure cycles) under basal conditions and after sequential injection of oligomycin (1 µM; OM), fluoro-carbonyl cyanide phenylhydrazone (1 µM; FCCP), and antimycin A (1 µM; AA) plus rotenone (1 µmol/L; ROT; all from Sigma-Aldrich).

### Human carotid artery plaques

Human carotid artery plaques were harvested during carotid artery endarterectomy (CEA) surgery, transported to the laboratory and snap frozen. CEA was performed due to advanced atherosclerotic lesion formation and stenosis in the carotid arteries. The patients’ characteristics are summarized in Supplemental Table 6. The tissue homogenization was performed in 700 µl Qiazol lysis reagent and total RNA was isolated using the miRNeasy Mini Kit (Qiagen, Netherlands) according to manufactureŕs instruction. RNA concentration and purity were assessed using NanoDrop. RIN number was assessed using the RNA Screen Tape (Agilent, USA) in the Agilent TapeStation 4200. Normalized gene expression levels obtained by RNA sequencing were used for linear regression analysis using the “ggcorrplot” R package(v0.1.4). A two-tailed P<0.05 was considered to be statistically significant.

## Statistics

Statistical analyses were performed using GraphPad Prism version 8.0 or higher (GraphPad Software, Inc.). Outliers were identified with ROUT = 1 and normality of the data was tested via the D’Agostino–Pearson omnibus normality test. To compare 2 normally distributed groups of data without differences in the variances, an unpaired Student t test was performed. For >2 groups, the data were compared by one-way or two-way ANOVA. For post hoc analysis, the Sidak, Tukey, or Dunett test was performed to correct for multiple comparisons and the Fisher Least Significant Difference test was used for planned comparisons. All data are shown as mean ± s.e.m. A two-tailed P < 0.05 was considered statistically significant.

## Study approval

All animal experiments were approved by the local Ethics committee (District Government of Upper Bavaria; License Number: 55.2-1-54-2532-111-13 and 55.2-2532.Vet_02-18-114) and conducted in accordance with the institutional and national guidelines and following the ARRIVE guidelines. Human patients provided written informed consent in accordance with the Declaration of Helsinki. The study has been approved by the local Ethics Committee of the Klinikum rechts der Isar of the Technical University Munich (diary numbers 2799/10 and 2799/10S).

## Author contributions

Y.W. carried out experiments, analyzed data and drafted Figures; G.L. performed bioinformatic RNAseq data analyses; B.C., G.S. and M.V. contributed to the *in vivo* experiments and data analysis; E.v.d.V. and S.M. provided the kinase activity data; C.M. and A.B. contributed to the extracellular flux analysis; M.N.J. contributed to the plaque LCM; Z.L., N.S. and L.M. provided human plaque data; M.H. performed flow cytometric cell sorting, M.L. provided support and advice for RNAseq experiments and bioinformatic analyses; B.L. provided floxed *Cnr1* mice; C.W. provided critical infrastructure and funding; S.H., R.G.P. and S.S. designed and supervised the study and planned experiments; R.G.P. and S.S. reviewed data and wrote the manuscript. All authors reviewed, edited and approved the manuscript.

## Acknowledgements

We are grateful to Seray Anak, Martina Geiger, Yvonne Jansen, and Soloyamaa Bayasgalan for excellent support with histological analyses and genotyping, and to the entire ZVH animal facility team for their continuous support.

---

**Supplemental Figure 1.**
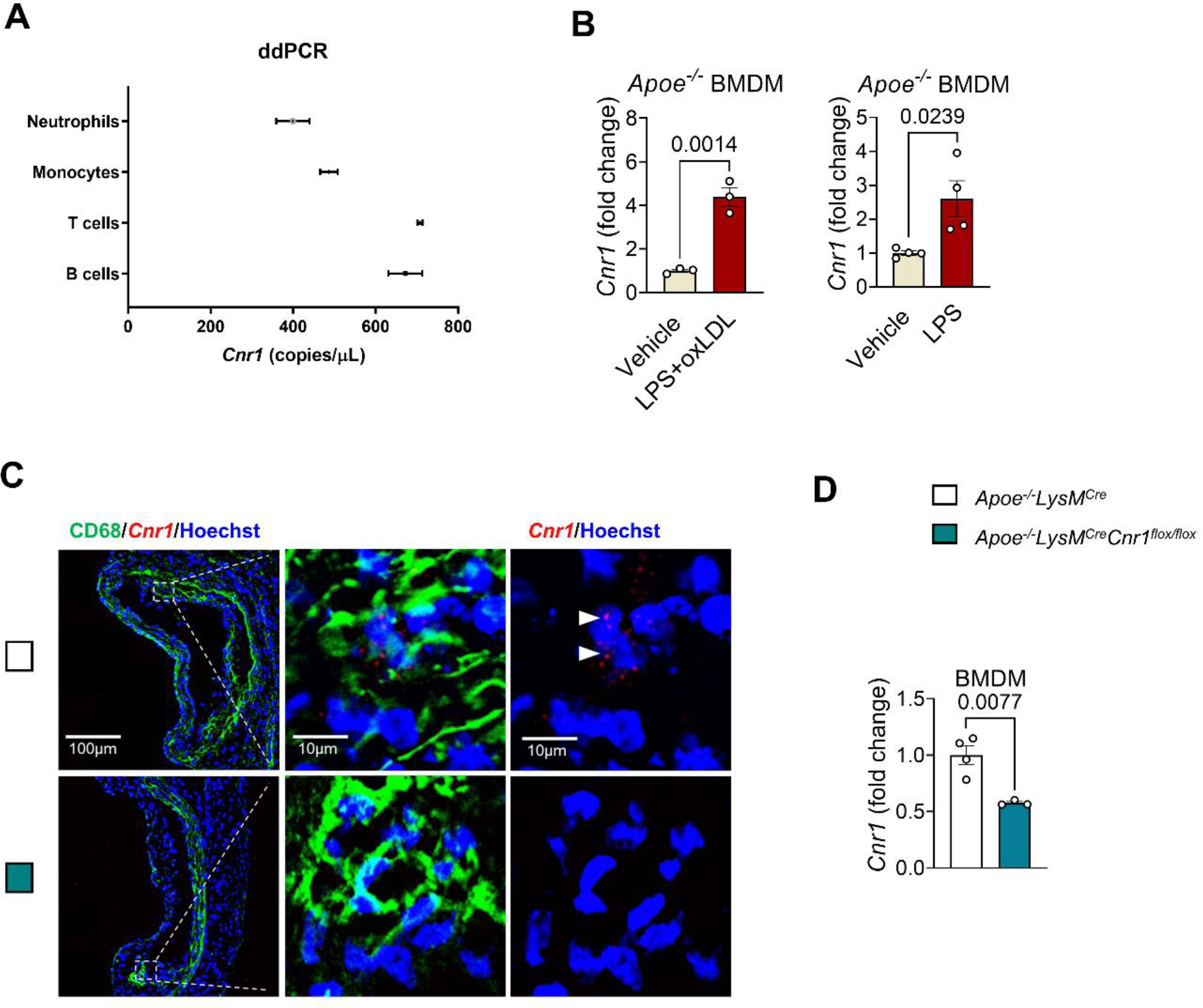
Detection of murine *Cnr1* expression in myeloid cell subsets and atherosclerotic plaques. (**A**) ddPCR analysis of *Cnr1* expression in sorted myeloid cell subsets (n=3). (**B**) qPCR analysis of *Cnr1* expression in vehicle-, LPS-(10 ng/mL) or oxLDL (1 µg/ml)-treated BMDMs from *Apoe^-/-^* male mice (n=3-4). (**C**) Fluorescence *in situ* hybridization of *Cnr1* co-immunostained for CD68 (green) and Hoechst33342 (nuclei, blue) in aortic root sections of *Apoe^-/-^* mice subjected to 4 weeks WD. Scale bar: 100 μm (left) and 10 µm (right). (**D**) RT-PCR analysis of *Cnr1* expression in BMDMs from *Apoe^-/-^LysM^Cre^*and compared with *Apoe^-/-^LysM^Cre^Cnr1^flox/flox^* mice (n=3-4). Each dot represents one mouse and all data are shown as mean ± s.e.m. Two-sided unpaired Student’s t-test (**B** and **D**) was used to determine the *P* value.

**Supplemental Figure 2.**
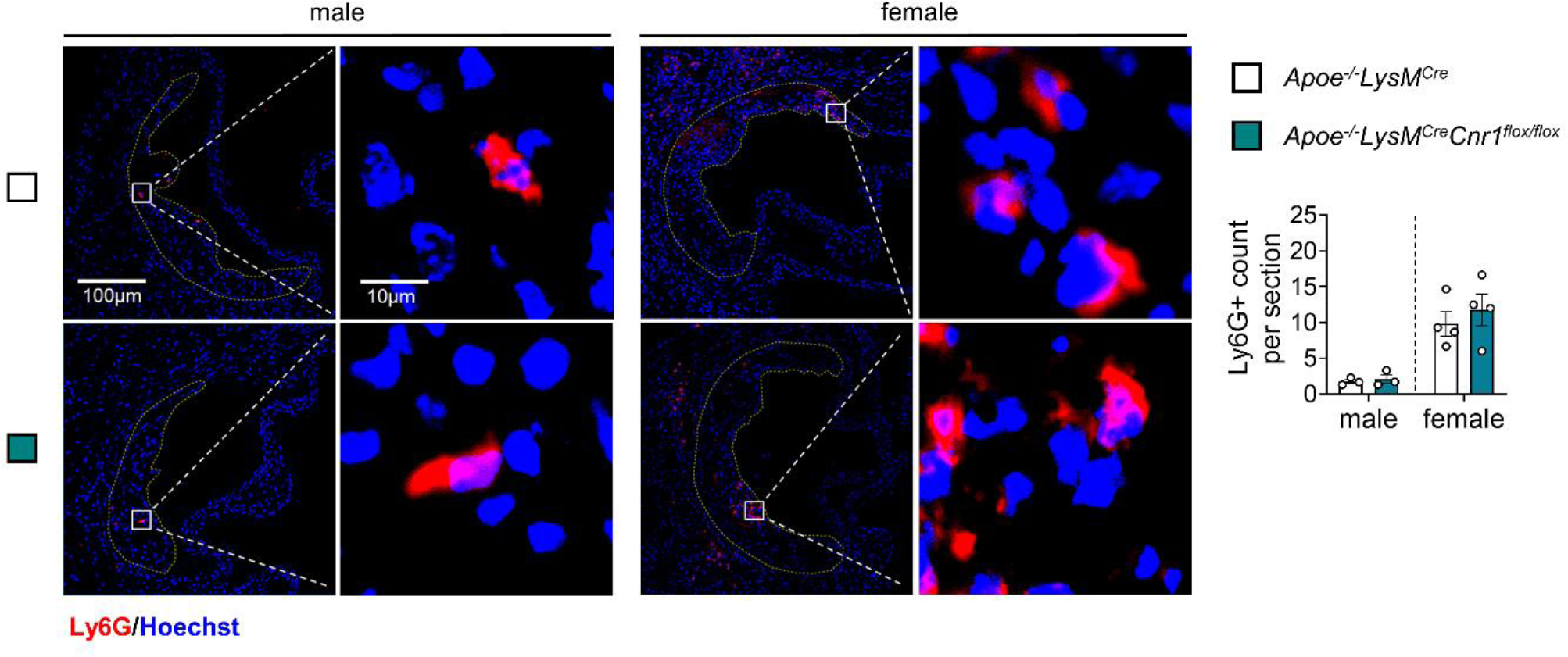
Impact of myeloid *Cnr1* deficiency on neutrophil recruitment at early stage of atherosclerosis. Representative immunostainings and quantification of neutrophils (Ly6G, red) in aortic root sections of male and female *Apoe^-/-^LysM^Cre^*and *Apoe^-/-^ LysM^Cre^Cnr1^flox/flox^* mice (n=3-4) after 4 weeks of WD; nuclei are counterstained with Hoechst33342 (blue). Scale bar: 100 μm (left) and 10 µm (right). Each dot represents one mouse and all data are expressed as mean ± s.e.m. Two-sided unpaired Student’s t-test. Male and female were analyzed independently.

**Supplemental Figure 3.**
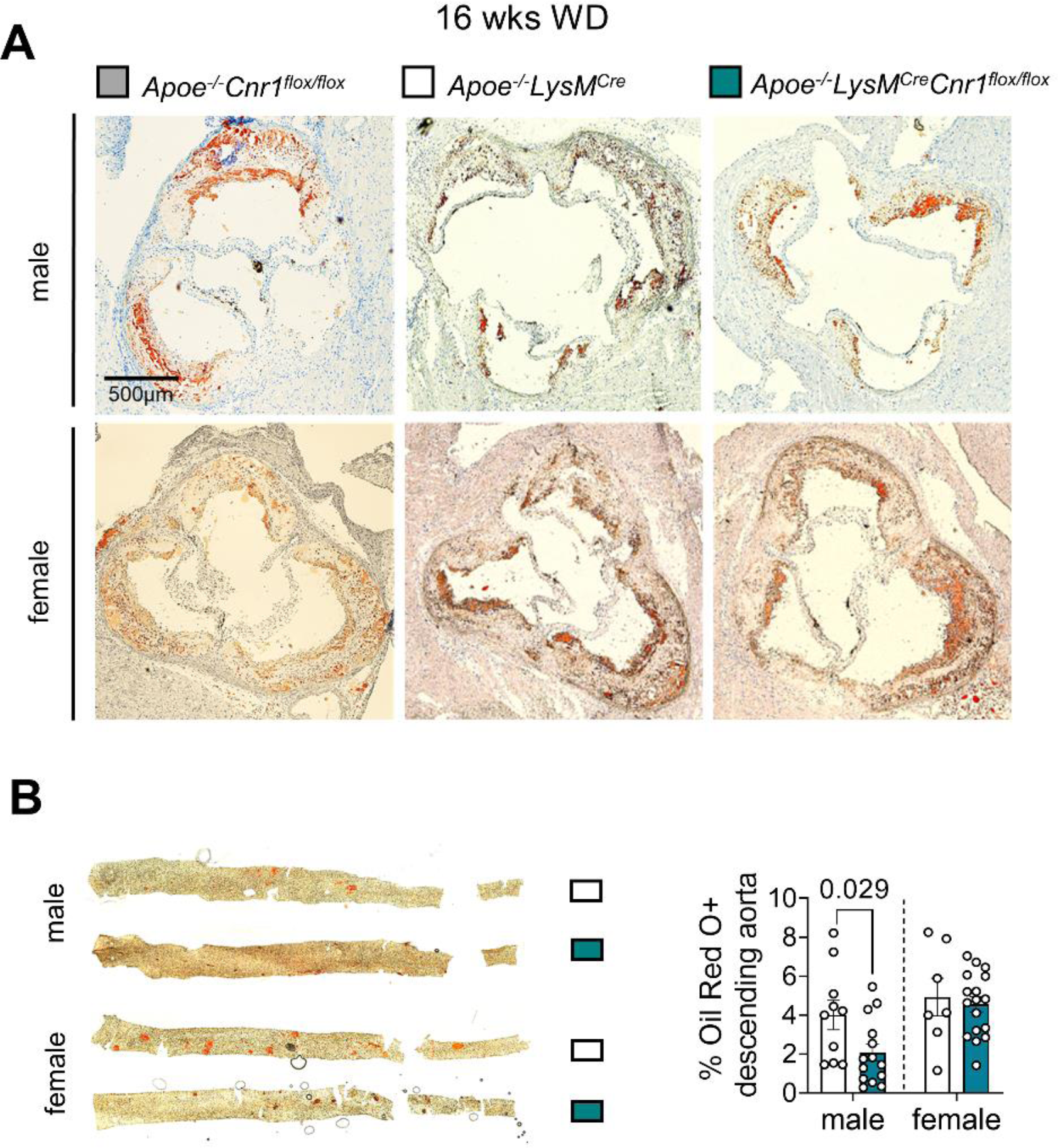
Impact of myeloid *Cnr1* deficiency on advanced stage of atherosclerosis. (**A**) Representative Oil-Red-O stains of aortic roots of male and female *Apoe^- /-^Cnr1^flox/flox^, Apoe^-/-^LysM^Cre^* and *Apoe^-/-^LysM^Cre^Cnr1^flox/flox^* mice after 16 weeks of WD. Scale bar: 500 μm. (**B**) Plaque area within descending thoraco-abdominal aortas after 16 weeks WD determined by en face preparations stained with Oil-red-O, and calculated as percentage of plaque per total vessel area (n=7-17). Each dot represents one mouse and all data are expressed as mean ± s.e.m. Two-sided unpaired Student’s t-test (**B male**) or Welch’s t test (**B female**). Male and female were analyzed independently.

**Supplemental Figure 4.**
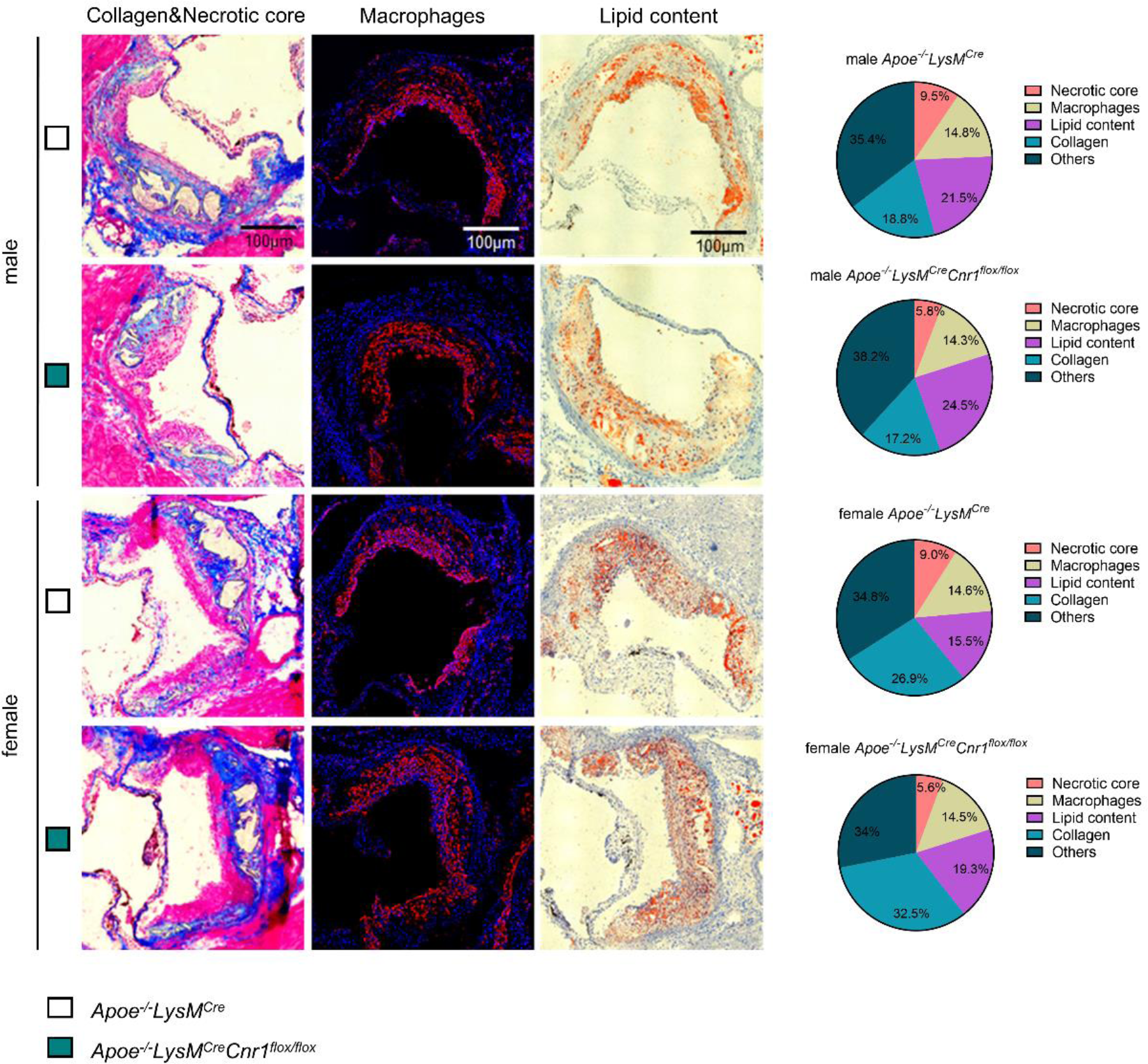
Impact of myeloid *Cnr1* deficiency on advanced plaque composition. Representative aortic root plaque images stained with Oil-red-O and Masson staining’s for lipid or collagen content, respectively, and quantification of the necrotic core area after 16 weeks WD. Scale bar: 100 μm. Necrotic core and collagen content per total plaque area (n=6-10) were calculated. The accumulation of lesional macrophages was determined by CD68 staining (n=6-10).

**Supplemental Figure 5.**
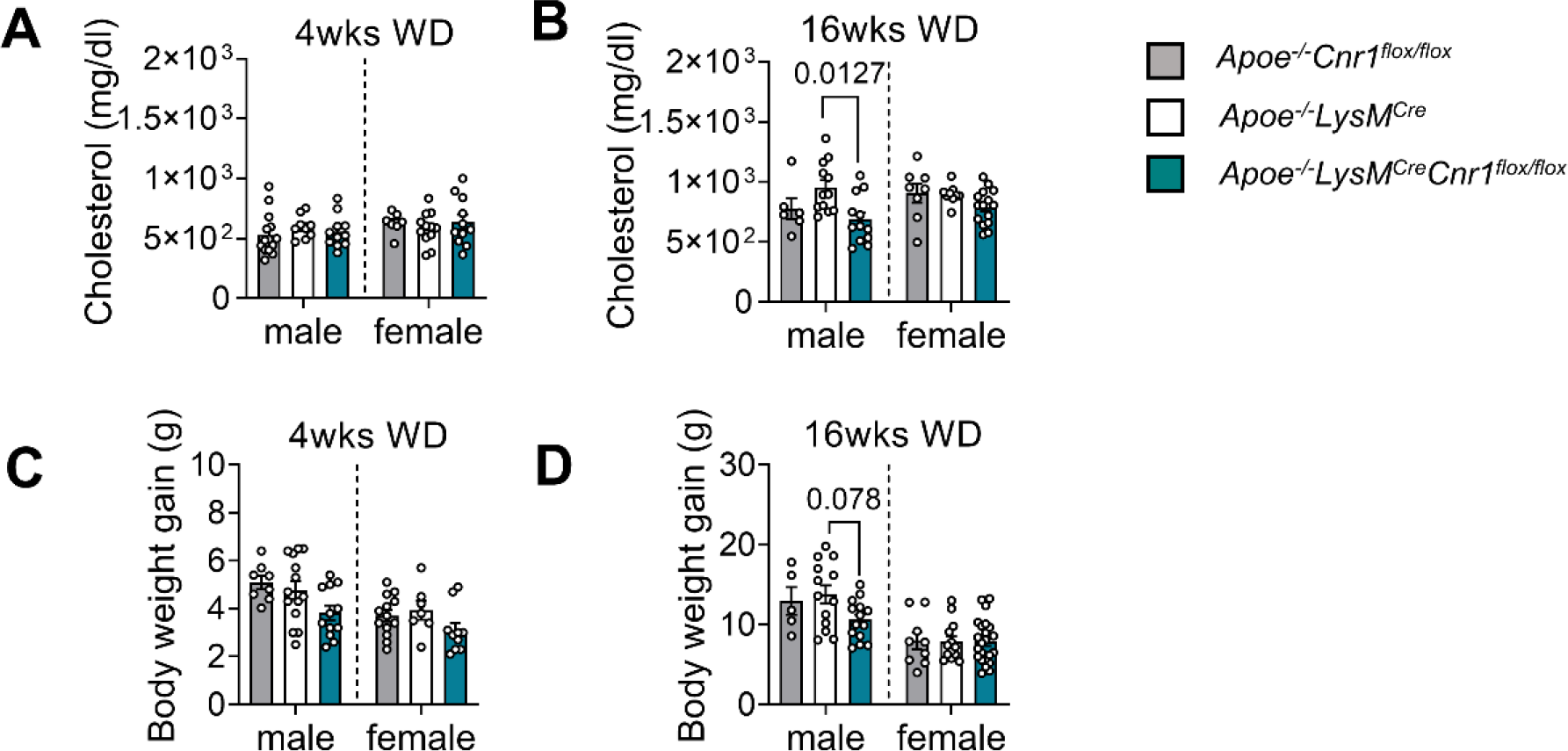
Impact of myeloid *Cnr1* deficiency on metabolic parameters. (**A-B**) Plasma cholesterol levels after 4 and 16 weeks Western diet (WD) in *Apoe^-/-^Cnr1^flox/flox^, Apoe^-/-^LysM^Cre^* and *Apoe^-/-^LysM^Cre^Cnr1^flox/flox^* mice (n=6-14). (**C-D**) Body weight gain after 4 and 16 weeks WD respect to the initial body weight in *Apoe^-/-^Cnr1^flox/flox^, Apoe^-/-^LysM^Cre^* and *Apoe^-/-^ LysM^Cre^Cnr1^flox/flox^*mice (n=5-23). Each dot represents one mouse and all data are expressed as mean ± s.e.m. one-way ANOVA followed by Dunnett T3 test (**A-D**). Male and female were analyzed independently.

**Supplemental Figure 6.**
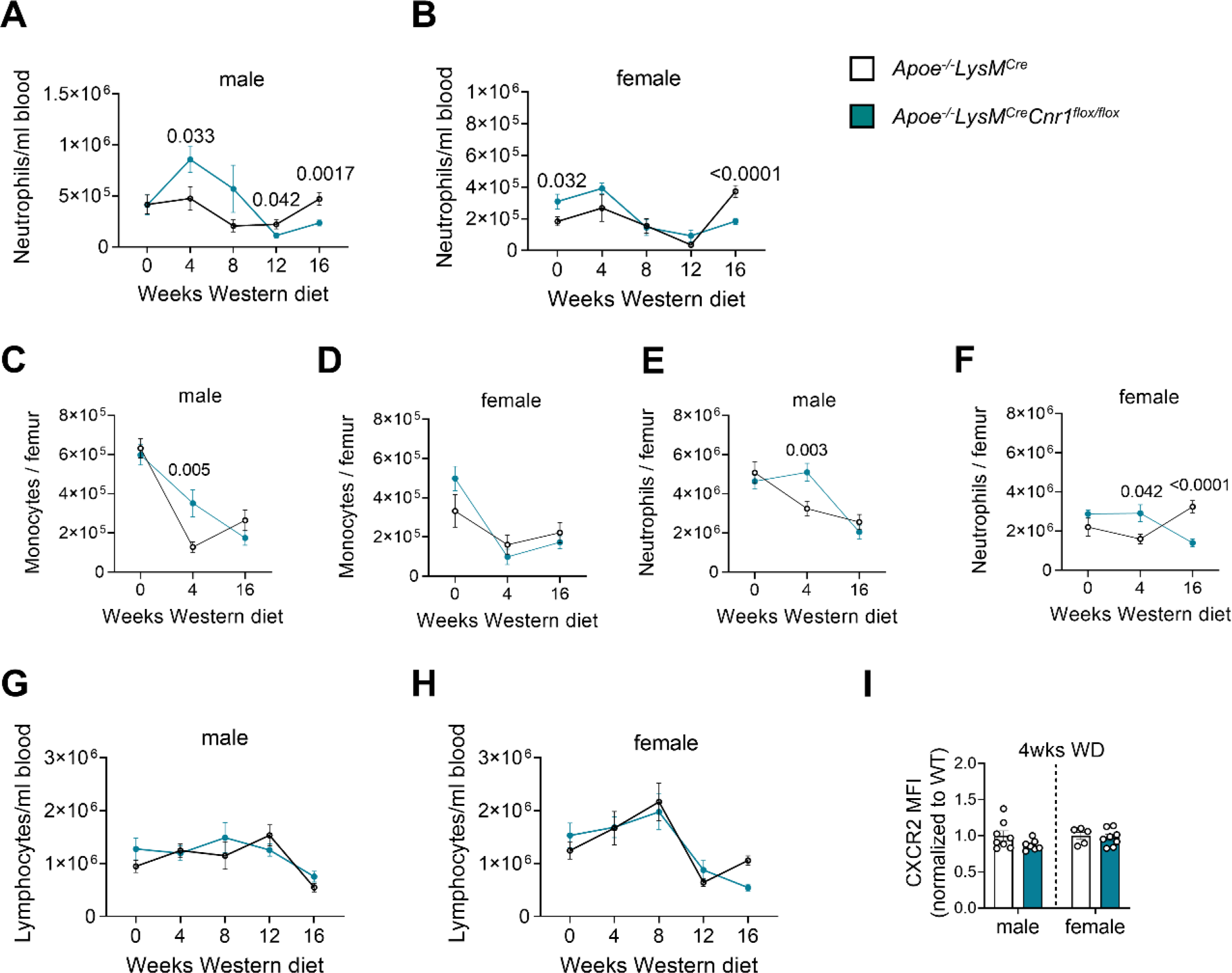
Impact of myeloid *Cnr1* deficiency on leukocyte subsets counts. Number of circulating neutrophils in male (m) and female (f) mice assessed by flow cytometry after 0 (n=9-10), 4 (n=18-19 m; n=6-10 f) 8 (n=10 m; n=5-10 f), 12 (n=7-8 m; n=5-6 f) and 16 (n=14-15 m; n=15-23 f) weeks of Western diet (WD) (**A** and **B**). Number of femural monocytes gated as CD45+ CD11b+ CD115+ (**C** and **D**) and neutrophils gated as CD45+ CD11b+ Ly6G+ (**E** and **F**) assessed by flow cytometry after 0, 4 and 16 weeks of WD. Number of circulating lymphocytes assessed by flow cytometry after 0, 4, 8, 12 and 16 weeks of WD (**G** and **H**). **I**, Flow cytometric analysis of CXCR2 surface expression on circulating neutrophils after 4 weeks of WD (n=5–8). Each dot represents one mouse and all data are expressed as mean ± s.e.m. Two-sided unpaired Student’s t-test (**A-I**). Male and female were analyzed independently.

**Supplemental Figure 7.**
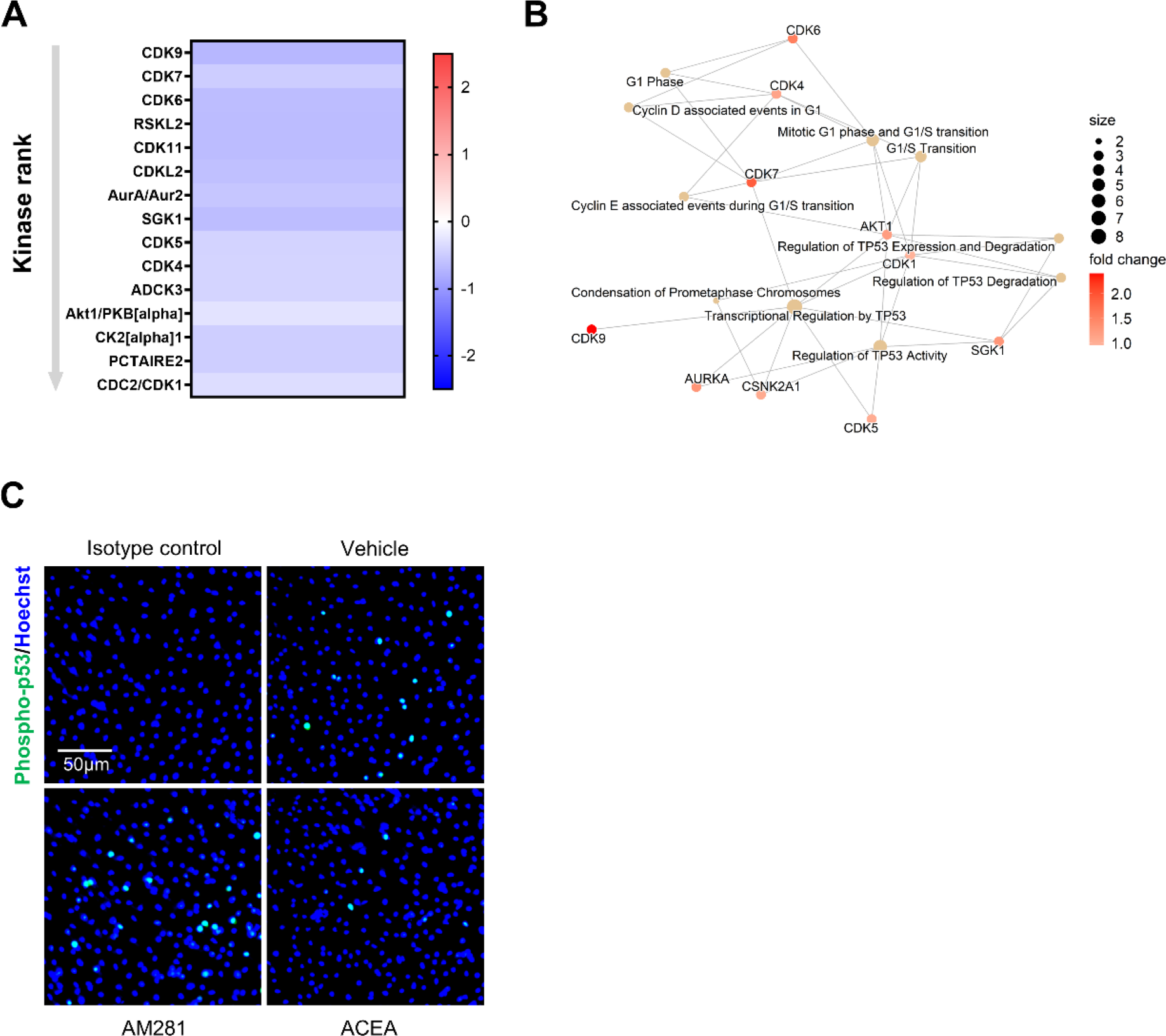
Effect of CB1 stimulation on kinase activity and p53 nuclear translocation. (**A**) Heatmap of kinase activity in *Apoe^-/-^* BMDM treated with vehicle or 1 µM ACEA for 60 min. Values indicate mean kinase statistic; significantly different kinases with median final score >1.2 are shown. (**B**) Interaction map showing kinase pathways regulated in ACEA-stimulated BMDMs. (**C**) Phospho-p53 immunostaining (green) of *Apoe^-/-^* BMDMs treated with vehicle, 1 µM ACEA or 10 µM AM281 for 60 min. Nuclei were stained with Hoechst33342 (blue); scale bar: 50 μm.

**Supplemental Figure 8.**
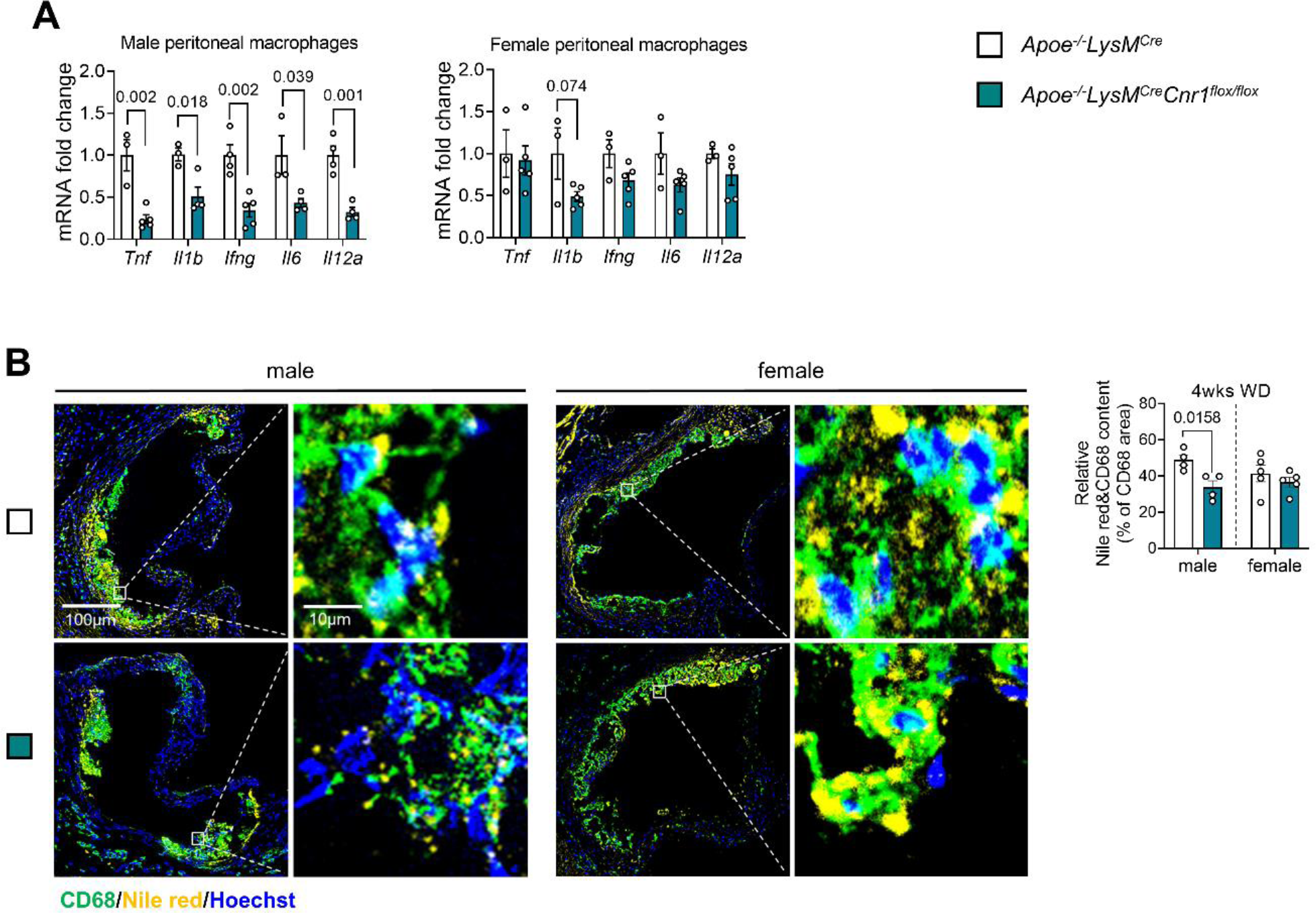
Impact of myeloid *Cnr1* deficiency on peritoneal macrophage inflammatory gene expression and plaque macrophage lipid droplet accumulation. **(A)** Expression levels of pro-inflammatory genes in peritoneal macrophages of *Apoe^-/-^LysM^Cre^* and *Apoe^-/-^LysM^Cre^Cnr1^flox/flox^*mice after 4 weeks Western diet (WD) (n=3-5). (B) Nile Red staining (yellow) of lipid droplets within aortic root plaque macrophages (CD68, green) after 4 weeks WD. Double immunostaining of Nile red and CD68 in aortic root lesions of male and female *Apoe^-/-^LysM^Cre^*and *Apoe^-/-^LysM^Cre^Cnr1^flox/flox^* mice after 4 weeks of WD (n=4-5). The accumulation of lipid droplets in macrophages were quantified by combined CD68 and Nile red staining. Nuclei were stained with Hoechst33342 (blue); Scale bar: 100 μm (left) and 10 µm (right). Each dot represents one mouse and all data are expressed as mean ± s.e.m. Two-sided unpaired Student’s t-test. Male and female were analyzed independently.

**Supplemental Figure 9.**
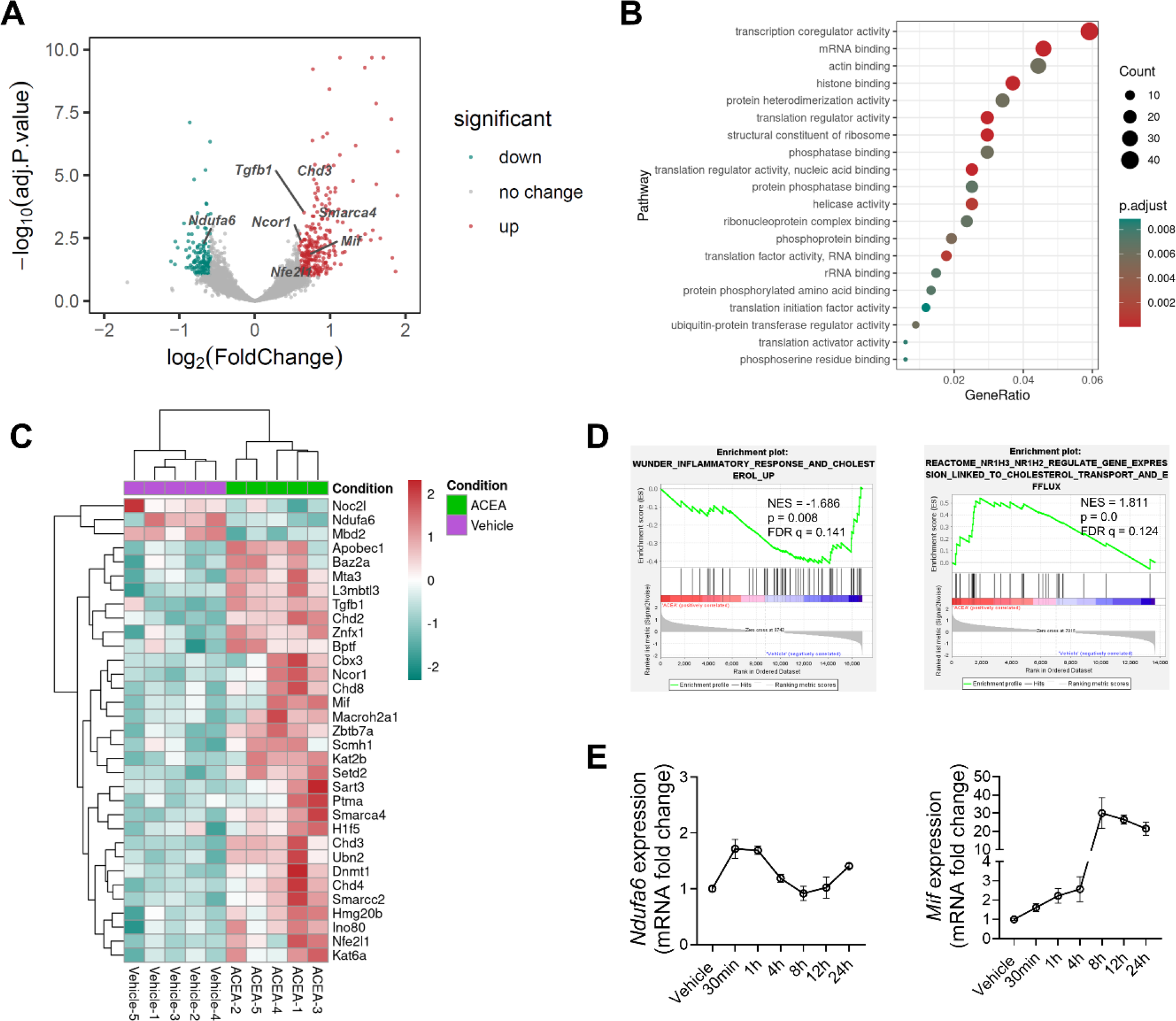
Impact of CB1 stimulation on the transcriptomic profile of BMDM. **(A)** Volcano plot of bulk RNA-seq data showing differentially expressed genes (DEGs) in male *Apoe^-/-^* BMDMs stimulated with 1 µM ACEA for 24 h versus vehicle (n=5 separate BMDM donor mice). (**B**) Pathway analysis showing top molecular functions regulated in CB1-stimulated BMDM. (**C**) Heat map showing chromatin modification pathway-related DEGs and selected genes highlighted in (**A**). (**D**) Gene set enrichment analysis (GSEA) identifies CB1-regulated gene sets associated with inflammatory response and cholesterol. Enrichment plots of significantly enriched gene sets with normalized enrichment score (NES), nominal P value and false discovery rate (FDR) q-value are shown. (**E**) qPCR analysis of male *Apoe^-/-^* BMDMs stimulated with 1 µM ACEA for the indicated time points, normalized to vehicle (time point 0). Each dot represents one mouse and all data are expressed as mean ± s.e.m. Two-sided unpaired Student’s t-test (**D**). Male and female were analyzed independently.

**Supplemental Figure 10.**
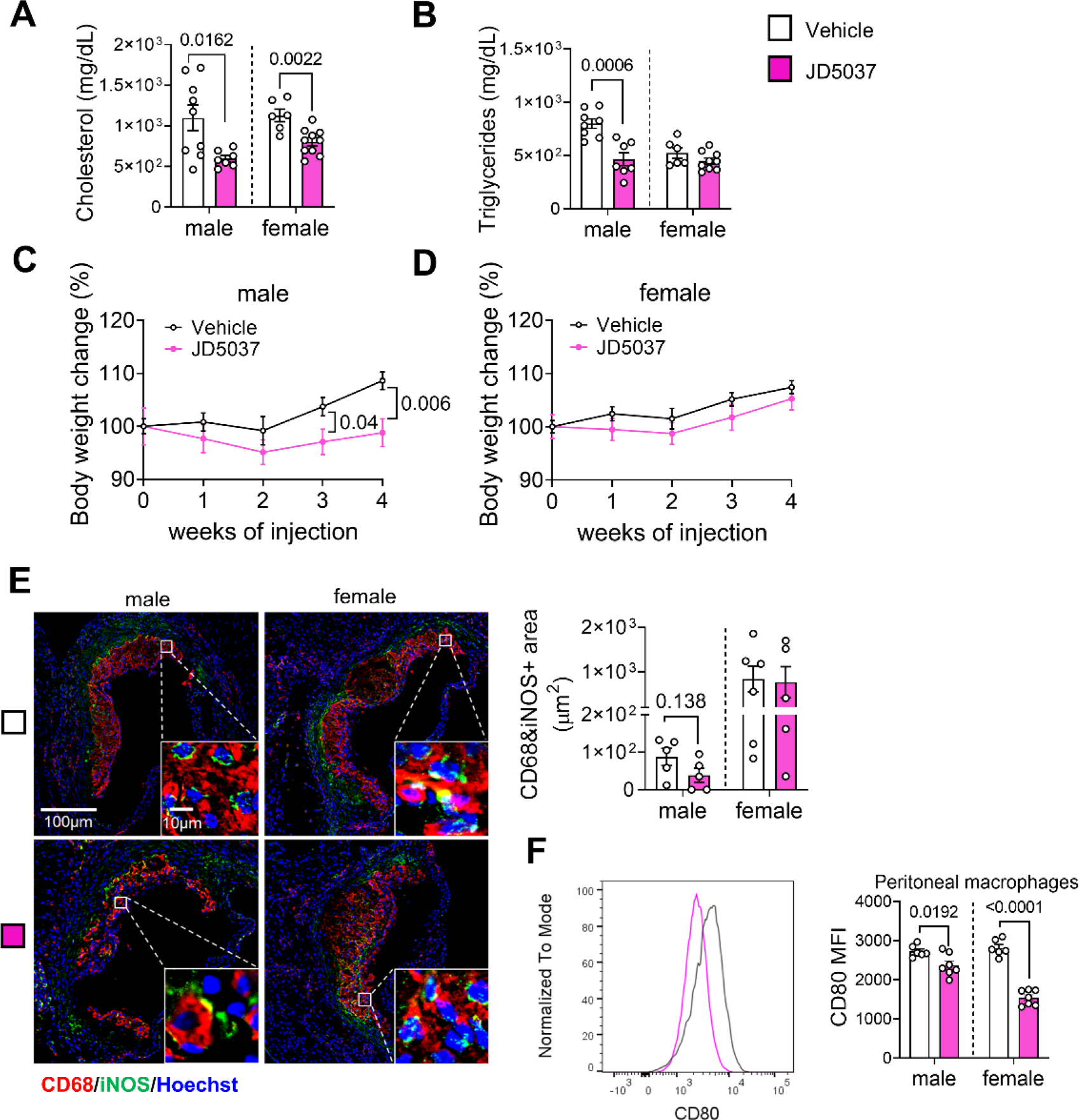
Impact of peripheral CB1 antagonist JD5037 on metabolic parameters, plaque macrophage accumulation and phenotype. **(A-B)** Plasma cholesterol and triglyceride levels and (**C-D**) body weights in male and female *Ldlr^-/-^* mice (n=6-10) after a total of 8 weeks WD diet, and receiving daily injections of vehicle- or peripheral CB1 antagonist JD5037 for the last 4 weeks of diet. (**E**) Double immunostaining and quantification of iNOS (green) and CD68 (red)-positive macrophages in aortic root lesions of male and female *Ldlr^-/-^* mice (n=5-6) treated with vehicle or JD5037. Nuclei were stained with Hoechst33342 (blue). Scale bar: 100 μm (left) and 10 µm (right). (**F**) Surface expression of the pro-inflammatory macrophage marker CD80 on peritoneal macrophages, assessed at the study endpoint (n=6-7). Each dot represents one mouse and all data are expressed as mean ± s.e.m. Two-sided unpaired Student’s t-test (**A-F**). Male and female were analyzed independently.

## Supplemental Material

Loss of myeloid cannabinoid receptor CB1 confers atheroprotection by reducing macrophage proliferation and immunometabolic reprogramming (Wang et al.)

**Supplemental Table 1:** Antibodies used for immunostaining.

| Antigen | Source | Dilution | Reference | Provider |
| --- | --- | --- | --- | --- |
| CD68 | Rat | 1:400 | MCA1957GA | Bio-Rad |
| Ki67 | Rat | 1:50 | 11-5698-82 | eBioscience |
| Ly6G | Rat | 1:100 | 551459 | BD |
| NOS(pan) | Rabbit | 1:100 | 2977 | Cell Signaling |
| Phospho-p53 | Rabbit | 1:100 | 12571s | Cell Signaling |

**Supplemental Table 2:** Isotype controls.

| Immunoglobulin | Reference | Provider |
| --- | --- | --- |
| Normal rat IgG | 6-001-A | RD system |
| Normal rabbit IgG | 315-005-003 | JIR |
| Rat IgG2a FITC | 553929 | BD |

The working concentration of normal IgG for isotype control was same to specific primary antibody. IgG = Immunoglobulin G; JIR = Jackson ImmunoResearch.

**Supplemental Table 3:** Secondary antibodies.

| Antigen | Source | Conjugation | Dilution | Reference | Provider |
| --- | --- | --- | --- | --- | --- |
| anti-rabbit | donkey | AF488 | 1:100 | 711-545-152 | JIR |
| anti-rat | donkey | AF488 | 1:300 | 712-546-150 | JIR |
| anti-rat | donkey | Cy3 | 1:300 | 712-165-153 | JIR |

**Supplemental Table 4:**
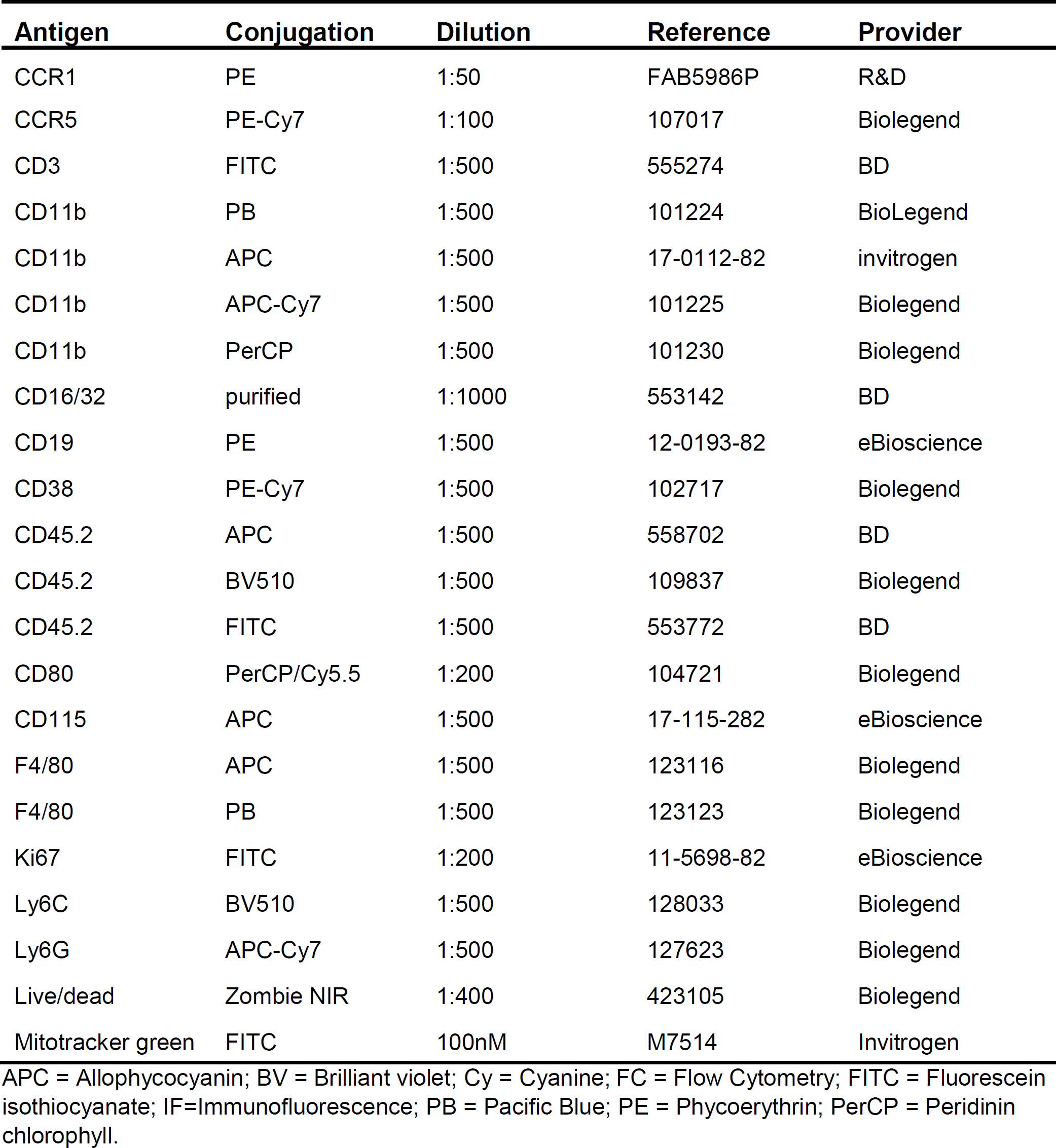
Murine FACS antibodies.

**Supplemental Table 5:**
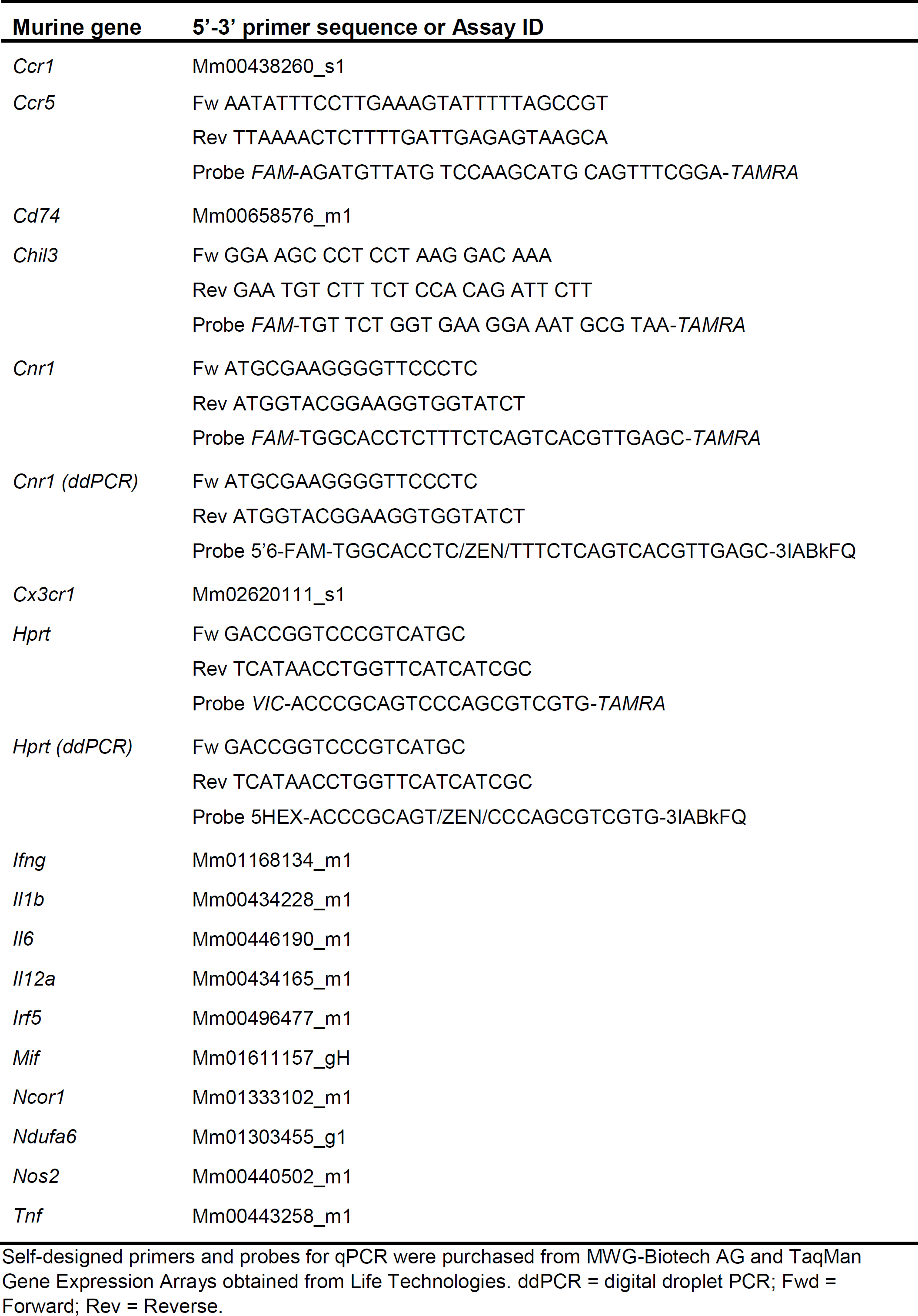
Primers for qPCR and ddPCR analysis.

**Supplemental table 6:** Patient characteristics.

| Patient characteristics |  | Total Number (n= 171) |
| --- | --- | --- |
| Age (mean, range), years |  | 72 (53-93) |
| Gender | male | 121 (70,8 %) |
|  | female | 50 (29,2 %) |
| BMI (Mean) |  | 26,8 |
| ASA Stadium (Mean) |  | 2,5 |
| Hypertension |  | 130 (78,3 %)* |
| CHD |  | 39 (22,8 %) |
| Diabetes |  | 37 (22,3 %) * |
| Dyslipidemia |  | 99 (59,6 %)* |
| Dialysis |  | 0 |
| Smoking | active smoker | 23 (13,9 %) |
|  | ex smoker | 47 (28,5 %) |
|  | never | 95 (57,6%) |
|  | n/a | 6 |
| State at presentation | symptomatic | 69 (40,4 %) |
|  | asymptomatic | 102 (59,6 %) |
| Type of symptoms | TIA | 4 (5,8 %) |
|  | ipsilateral hemispheric TIA | 22 (31,9 %) |
|  | amaurosis fugax | 2 (2,9 %) |
|  | amaurosis fugax ipsilateral | 21 (30,4 %) |
|  | apoplex | 4 (5,8 %) |
|  | ischemic stroke before elective surgery | 14 (20,3 %) |
|  | acute/progressive stroke | 2 (2,9 %) |
| Degree of stenosis<br>(mean, range), % |  | 81,5 (40-100) |
| TIA, transient ischemic attack |  |  |

